# Spatiotemporal single-cell RNA sequencing of developing hearts reveals interplay between cellular differentiation and morphogenesis

**DOI:** 10.1101/2020.05.03.065102

**Authors:** Madhav Mantri, Gaetano J. Scuderi, Roozbeh Abedini Nassab, Michael F.Z. Wang, David McKellar, Jonathan T. Butcher, Iwijn De Vlaminck

**Author notes:** These authors contributed equally (Madhav Mantri, Gaetano J. Scuderi).

## Abstract

Single-cell RNA sequencing is a powerful tool to study developmental biology but does not preserve spatial information about cellular interactions and tissue morphology. Here, we combined single-cell and spatial transcriptomics with new algorithms for data integration to study the early development of the chicken heart. We collected data from four key ventricular development stages, ranging from the early chamber formation stage to the late four-chambered stage. We created an atlas of the diverse cellular lineages in developing hearts, their spatial organization, and their interactions during development. Spatial mapping of differentiation transitions revealed the intricate interplay between cellular differentiation and morphogenesis in cardiac cellular lineages. Using spatially resolved expression analysis, we identified anatomically restricted gene expression programs. Last, we discovered a stage-dependent role for the small secreted peptide, thymosin beta-4, in the coordination of multi-lineage cellular populations. Overall, our study identifies key stage-specific regulatory programs that govern cardiac development.

## INTRODUCTION

The heart is the first fully functional organ to develop and is vital for embryogenesis[1]. Cardiogenesis involves heterogeneous cell populations from multiple lineages that spatiotemporally interact to drive cardiac fate decisions[2]. The heterogeneity of cell types during cardiac development makes it difficult to study cardiac fate decisions using traditional developmental biology techniques. Single-cell RNA-sequencing (scRNA-seq) has been used to study the cellular mechanisms involved in driving heart development, but does not preserve spatial information, and does not enable studies of the complex interplay between cellular maturation and morphogenesis. Here, we combined spatially resolved RNA-seq with high-throughput scRNA-seq to study the spatiotemporal interactions and regulatory programs that drive fetal development of the chicken heart. Current spatial transcriptomics approaches lack single-cell resolution, which we addressed here using new approaches to integrate high-throughput spatial and single-cell transcriptomic data.

We used the chicken embryo as a model system to study cardiogenesis since the development of the chick ex-utero in an egg allows unique access to early stages of development when the heart consists of relatively few cells. The mature chick heart comprises four chambers with in- and out-flow tracts, and despite some differences, the chick heart anatomy resembles the anatomy of the human heart more closely than other non-mammalian vertebrate model organisms[3]. We generated over 22,000 single-cell transcriptomes across four key Hamburger-Hamilton ventricular development stages (HH21-HH24, HH30-HH31, HH35-HH36, and HH40). The data encompass common and rare cell types, including progenitor and mature cell types from multiple lineages. In addition, we performed spatially resolved RNA-seq on a total of 12 heart tissue sections collected at the same four stages.

As we demonstrate here, the combination of single-cell and spatial transcriptomics uniquely enables to unravel cellular interactions that drive cardiogenesis. The data enabled us to reconstruct a high-resolution, spatially resolved gene expression atlas of epi-, endo-, and myocardial developmental lineages within cardiac tissue. We characterized and spatially resolved progenitor and differentiated cell types, identified stage-specific transcriptional programs and cellular interactions, reconstructed differentiation lineages, and delineated important regulatory programs in cardiac development. We integrated scRNA-seq and spatial RNA-seq data using an anchor-based method to predict cell type annotations for spatially resolved transcriptomes. We used the cell-type predictions to construct proximity maps revealing novel cellular interactions. Using the cell-type prediction scores, we uncovered local cellular heterogeneity and spatially restricted regulatory programs in ventricular tissue and characterized changes in cellular environments across ventricular spatial compartments. We furthermore constructed a similarity map between single-cell and spatial transcriptomes, which enabled us to spatially map lineage-associated differentiation trajectories within the tissue. This analysis revealed spatiotemporal differentiation transitions within the epicardial lineage, and points to the utility of spatiotemporal single-cell RNA sequencing as a tool to study the interplay between cellular development and morphogenesis. Last, our analysis revealed a developmental stage dependent role for the small secreted peptide thymosin beta-4 in the coordination of heterogeneous multi-lineage cell populations across ventricular development stages and clarified its role in ventricular compaction and maturation.

## RESULTS

### Spatially resolved single-cell transcriptomics atlas of developing fetal chicken hearts

To study the complex interplay between differentiation and morphogenesis during cardiac development, we combined single-cell and spatial transcriptomics. We profiled four key Hamburger-Hamilton ventricular development stages of the chicken heart: ***i***) day 4 (HH21-HH24, whole ventricles), corresponding to the early chamber formation stage during the initiation of ventricular septation and only trabeculated myocardium, ***ii***) day 7 (HH30-HH31, left and right ventricles), one of the earliest stages of cardiac four chamber formation with ventricular septation almost complete but the myocardium containing mostly fenestrated trabeculated sheets, ***iii***) day 10 (HH35-HH36, left and right ventricles), a mid-four chamber development stage with septation complete and mostly compact myocardium, and ***iv***) day 14 (~HH40, left and right ventricles), a late four chamber development stage with completely compact myocardium and the main events of cardiogenesis essentially complete[3].

To perform scRNA-seq (10x Genomics chromium platform), we enzymatically digested cardiac ventricle tissue into single-cell suspensions (**Fig. 1A**, Methods). Pooling of cells from up to 72 fetal hearts enabled scRNA-seq on cardiac tissue at early stages of development. To perform spatial transcriptomics (10X Genomics Visium platform), we cryosectioned 10 ⎕m coronal tissue slices (chamber view) from fetal hearts at the same four stages (**Fig. 1A**, See Methods). We generated single-cell transcriptomic data for 22,315 cells and spatial transcriptomics data for 12 tissue sections covering over 6,800 barcoded spots (**Sup. Fig. 1A and 1B**). We found that spatial transcriptomes collected at the same developmental stage were strongly correlated (Pearson correlation; R > 0.98, **Sup. Fig. 1D**), and that spatial transcriptomes and single-cell transcriptomes collected at the same developmental stage were strongly correlated (Pearson correlation; R 0.88-0.91, **Sup. Fig. 1E**). The combination of scRNA-seq and spatial transcriptomics uniquely enabled us to spatially resolve cell-type specific gene expression in cardiac tissue (see below).

**Figure 1:**
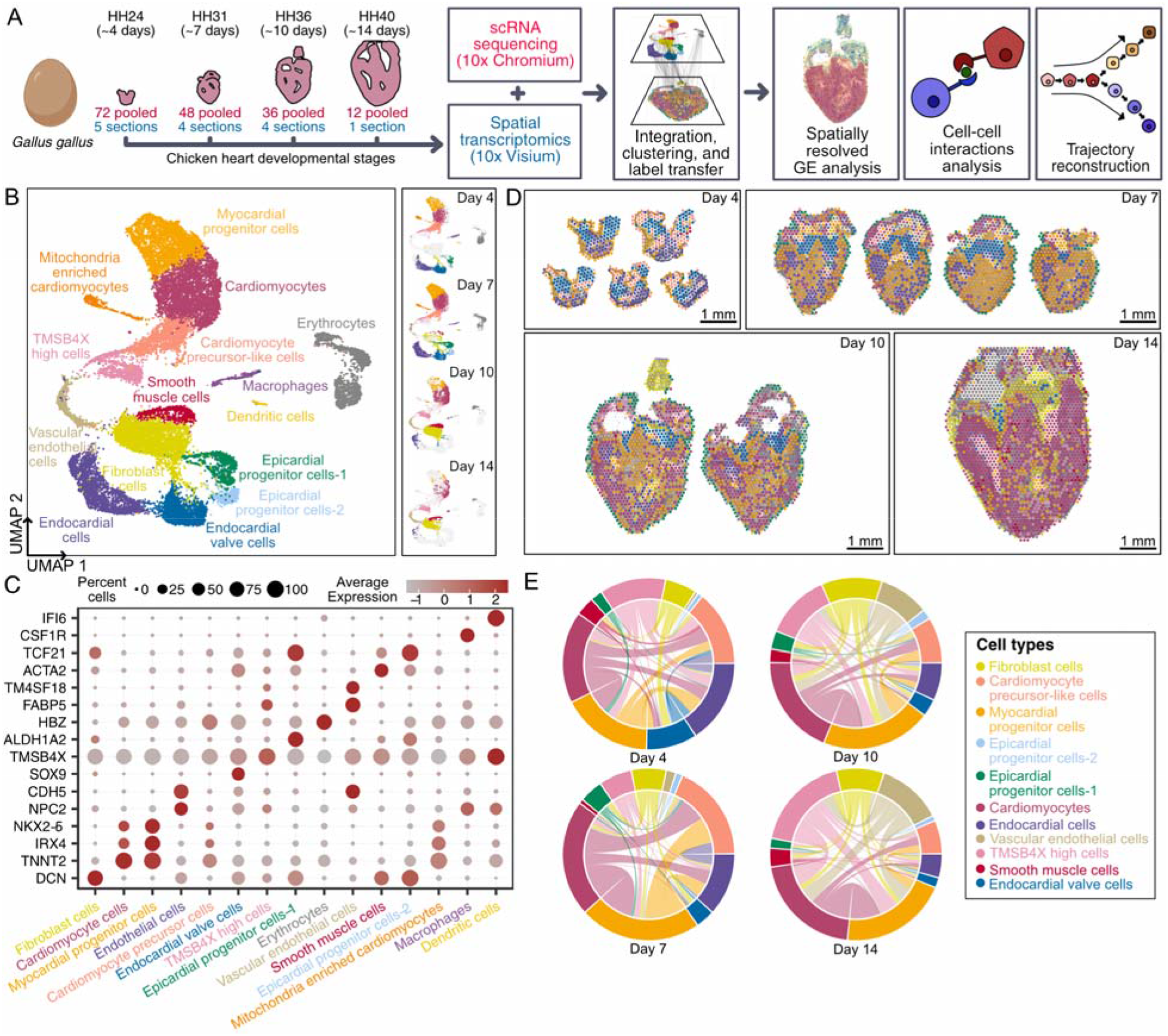
Spatially resolved single-cell transcriptomic atlas of developing fetal chicken hearts. **A)** Experimental workflow and analysis for single-cell RNA-seq and spatial RNA-seq of fetal chicken (Gallus *gallus*) hearts at four stages of development. **B)** UMAP projection of 22,315 single-cell transcriptomes clustered by gene expression and colored by cell type (Left). UMAP projection of single-cell transcriptomes colored by cell type and split by developmental stage from day 4 to day 14 (Right). **C)** Gene expression of cell type specific markers. Size of the dot represents the percent of cells in the clusters expressing the marker and the color intensity represents the average expression of the marker in that cluster. **D)** Spatial RNA-seq barcoded spots labeled by scRNA-seq cell type with maximum prediction score for four developmental stages. Only spots under tissue sections are shown. **E)** Chord diagrams representing cell-type proximity maps showing the degree of colocalization of cell type pairs within spots in spatial RNA-seq data across developmental stages.

To analyze single-cell transcriptomes, we filtered and preprocessed the data (Methods), performed batch correction using scanorama[4] (**Sup. Fig. 1C**), performed dimensionality reduction and cell clustering, and then visualized the data by Uniform Manifold Approximation and Projection (UMAP, Methods). This analysis revealed 15 distinct cell type clusters (**Fig. 1B**). We used marker gene and differential gene expression analysis to assign cell types to cell clusters (**Sup. Table 1**, **Fig. 1)**, and identified diverse cell clusters from myocardial, endocardial, and epicardial cardiac lineages in the ventricles (**Fig. 1C, Sup. Fig. 1F**). In addition to cardiac cell types, we detected a small number of circulating cell types including erythrocytes, macrophages, and dendritic cells. Last, we identified a unique heterogeneous population of cells that express high levels of thymosin beta-4 (TMSB4X, see below). A detailed overview of the cell-types identified is provided in the supplement (**Sup. Table 1**).

Standalone analysis of the spatial transcriptomic data revealed anatomical regions with differential transcriptional programs. To spatially resolve cell populations, the spatial transcriptomics data was integrated with the scRNA-seq data using Seurat-v3 anchor-based integration[5,6]. This approach first identifies anchors between datasets which represent pairwise correspondences between elements in the two datasets that appear to originate from the same biological state. The anchors are then used to harmonize the datasets by learning a joint structure with canonical correlation analysis and to transfer annotation information from one dataset to the other. Every spot in the spatial data could be considered a weighted mix of cell-types identified by scRNA-seq. We used the prediction scores from label transfer to obtain weights for each of the scRNA-seq-derived cell types for each spot (**Fig. 1D**, **Sup. Fig. 2**, Methods). In order to understand the spatial organization of cell types in broad anatomical regions, spots were labeled as cell types with maximum prediction score and visualized on H&E stained images of respective stages (**Fig. 1D**). Cell-type prediction scores for spatial transcriptomes were further used to estimate the abundance of pairs of specific cell types (Methods). As proximity is a necessity for physical interactions between two or more cells, these cell-type proximity maps can be used to guide the discovery of interactions between cell types from the same or different lineages. We constructed proximity maps for all cardiac cell type pairs and visualized them as chord diagrams (**Fig. 1E**). We found a significant number of cardiomyocytes colocalized with myocardial progenitor cells and precursor cells in all stages, as expected. We furthermore found a significant colocalization of myocardial cells with endocardial cells at day 7 and with vascular endothelial and fibroblast cells at day 10 and day 14. This was also expected given that endocardial cells line the trabeculated myocardium at day 7 and that vascular endothelial cells and fibroblasts are present in the compact myocardium by day 10.

### Spatially resolved cardiac lineage analysis

Ventricular development in fetal hearts involves regulatory interactions and the coordinated migration of cells from multiple lineages to form a fully developed four-chambered heart. We hypothesized that characterizing differentiation transitions in a spatial context would reveal cellular movements that occur during the differentiation process. To test this idea, we projected information about cellular transitions derived from scRNA-seq onto the spatial maps. We first gathered and reclustered single-cell transcriptomes from the epicardial, endocardial, and myocardial lineages **(Sup. Fig. 1F)**, and then reconstructed differentiation trajectories for these three lineages. Additional gene markers for these trajectory analyses can be found in the supplement (**Sup. Table 2**). We used PHATE[7] (Potential of Heat-diffusion for Affinity-based Transition Embedding) to visualize differentiation trajectories because of its ability to learn and conserve local and global structure in low dimensional space. We then spatially resolved sub-clusters of cell types and estimated the local PHATE1 dimension, a proxy for development time (**Methods, Fig. 2A**). Utilizing this approach, we found that the epicardial lineage undergoes a rich differentiation process while the endocardial and myocardial lineages mostly undergo a maturation process across the timepoints investigated here.

**Figure 2:**
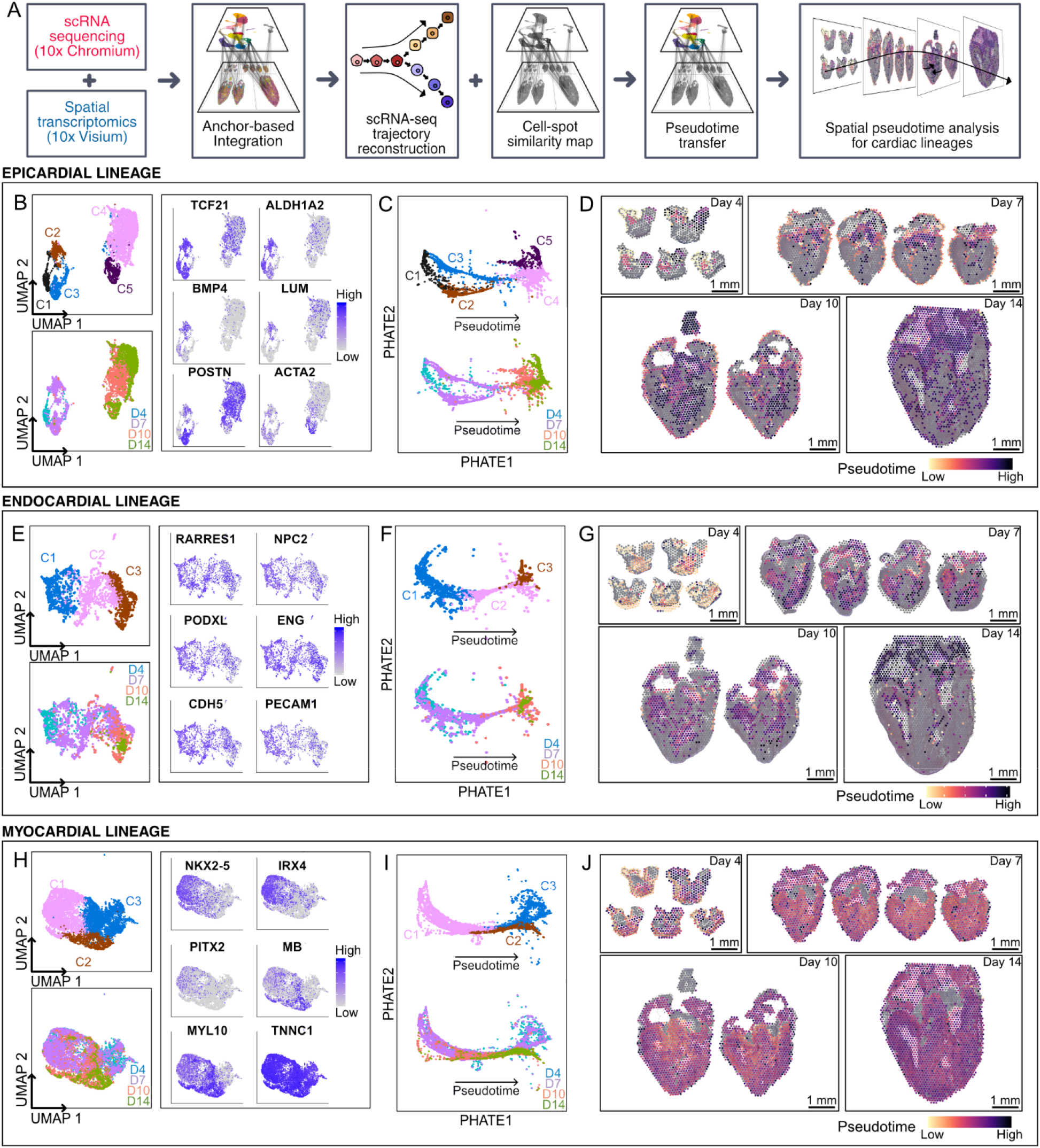
Spatiotemporal lineage analysis reveals heterogeneity in cardiac progenitor cells. **A)** Overview of analysis pipeline for trajectory reconstruction for scRNA-seq and spatial RNA-seq data. **B, E, H)** UMAP projection of single-cell transcriptomes from individual cardiac lineages clustered by gene expression and colored by cell type (left-top) and development stage (left-bottom). Feature plots showing expression of gene markers for cell types in individual cardiac lineages (right). **C, F, I)** Cardiac lineages visualized by PHATE and labeled by cell type (top) and development stage (bottom). **D, G, J)** Spatially resolved spatial RNA-seq spot pseudotime for cardiac lineage across developmental stages. Spot pseudotime was estimated using a similarity map between scRNA-seq cells and spatial RNA-seq spots. **B, C, D)** Epicardial lineage **E, F, G)** Endocardial lineage **H, I, J)** Myocardial lineage.

Lineage analysis for epicardial cells revealed a rich spatiotemporal differentiation process. The epicardial lineage was comprised of five cell clusters: an early epicardial progenitor cells cluster (C1), two intermediate precursor cell clusters (C2 and C3), a fibroblast cell cluster (C4), and a smooth muscle cell cluster (C5) (**Fig. 2B**, left). Cells in cluster C1 were mainly derived from the day 4 hearts and expressed the epicardial progenitor markers transcription factor 21 (TCF21) and t-box transcription factor 18[8] (TBX18), retinoic acid signaling-related transcripts aldehyde dehydrogenase[9,10] (ALDH1A2) and midkine[11,12] (MDK), as well as an epithelial-like phenotype marker keratin 7[13] (KRT7) (**Fig. 2B**, left and right). TCF21+ progenitor cells in cluster-C2 expressed bone morphogenetic protein 4 (BMP4), which is associated with an epicardial progenitor-like phenotype[14], and lumican (LUM), which is known to be expressed in the outermost epicardial layer of fetal hearts[15] (**Fig. 2B**, right). In contrast, TCF21+ cells in cluster-C3 expressed high levels of transcripts implicated in cell migration and differentiation, including fibronectin (FN1), periostin (POSTN), and agrin (AGRN)[16–19]. Cells in fibroblast cluster C4 expressed extracellular matrix markers, such as collagen type III (COL3A1) and periostin (POSTN). Smooth muscle cells in cluster C5 expressed ACTA2. PHATE-based trajectory reconstruction confirmed branching cell fates at day 7 and branch-merging and differentiation at day 10 (**Fig. 2C**). Transferring cell type labels to spatial data suggested that only the signaling epicardial progenitor cells (clusters C1 and C2) were spatially restricted to the outermost layer of the ventricular wall (**Sup. Fig. 2A**). Fibroblast and smooth muscle cells were found dispersed in the ventricular myocardium in later cardiac development stages (days 10 and 14, **Sup. Fig. 2C-D**). Pseudotime correlated well with stages in spatial data and revealed significant within-stage variability at days 7 and 10 with a presence of undifferentiated cells in the outer lining and differentiated cells in the myocardium (**Fig. 2D**). Additional lineage trajectory analyses by monocle-v2[20,21] are presented in the supplement (see Methods, **Sup. Fig. 3A-B**). Collectively, we find that at the day 4 stage, the epicardium is composed of signaling epicardial-progenitors that line the outermost layer of the ventricle. By day 7, the epicardial progenitors partition into two phenotypes: 1) an outermost epithelial-like layer that maintains a signaling progenitor phenotype and 2) a subepicardial layer that is actively differentiating and migrating into the myocardium with an upregulation of extracellular matrix directing cues. By day 10 and 14, the epicardial cells are fully differentiated into smooth muscle cells that are present in the myocardium and fibroblast-like cells that are present throughout the myocardium and line the epicardium. As expected, gene ontology terms of the epicardial lineage that significantly correlated with pseudotime revealed an upregulation of extracellular matrix related terms (**Sup. Fig. 3C**).

We further employed lineage analysis to study maturation of the endocardial and myocardial lineages. We identified three cell clusters within the endocardial lineage: early endocardial cells (C1) from days 4 and 7, intermediate endocardial cells (C2) from days 4 and 7, and terminally mature endocardial cells (C3) from days 10 and 14 (**Fig. 2E**, left). Cells from all three ventricular endocardial cell subclusters expressed the endocardial marker (NPC2), the differentiated endothelial markers cadherin (CDH5), podocalyxin[22] (PODXL), platelet and endothelial cell adhesion molecule 1[23] (PECAM1), and endoglin[24] (ENG), and the retinoic acid signaling related transcript RARRES1[25]. Endocardial cells lined the ventricular chambers of all heart stages in the spatial RNA-seq data (**Sup. Fig. 2E**). PHATE based trajectory analysis revealed endocardial cells undergo a maturation process from day 4 to day 14 and this was confirmed by independent trajectory analysis by Monocle v2 (**Fig. 2F-G & Sup. Fig. 3D-E**). In the myocardial lineage we identified three clusters, a myocardial progenitor cell cluster (C1) predominantly populated with cells from day 4 and 7 tissues, a differentiated cardiomyocyte cell cluster (C2) from mostly days 10 and 14, and a cardiomyocyte precursor-like cell cluster (C3) from days 4 and day 7 (**Fig. 2H, left**). All cells expressed cardiac troponin (TNNC1) while myocardial progenitor cells differentially expressed the progenitor markers NKX2-5[26], PITX2[27], and IRX4[28]. The cardiomyocytes were enriched in myosin light chain-10 (MYL10) and myoglobin (MB) (**Fig. 2H, right**). PHATE-based and monocle trajectory reconstruction of the myocardial lineage confirmed a differentiation path from myocardial progenitors (C1) to differentiated cardiomyocytes (C2), but the cardiomyocyte precursor-like cell cluster (C3) did not follow this trajectory (**Fig. 2I**& **Sup. Fig. 3G, 3H**). Transferring cell type labels on the spatial maps demonstrated that myocardial progenitor cells (C1) decrease in abundance with developmental stage while differentiated cardiomyocytes (C2) increase in abundance with developmental stage in the myocardium (**Sup. Fig. 2F-G**). The cardiomyocyte precursor-like cells (C3) were present in the myocardium across all stages (**Sup. Fig. 2H**). Overall, cells among cluster C3 remain an unknown cell cluster of cardiomyocytes. Pseudotime correlated well with stages in spatial data and confirmed simultaneous myocardial differentiation and myocyte maturation from day 4 to day 14 (**Fig. 2J**). Collectively, these results indicate that the endocardial and myocardial cells are a differentiated phenotype as early as the day 4 heart stage and mature with development time from day 4 to day 14. Gene ontology terms of the endocardial and myocardial lineage that significantly correlated with pseudotime can be found in supplement (**Sup. Fig. 3F-I**).

Transferring the valve endocardial cell type labels from scRNA-seq to spatial transcriptomics data confirmed that these cells are spatially restricted to atrioventricular heart valves at all four stages in the spatial data (**Sup. Fig. 2I**). Therefore, day 4 and day 7 valve endocardial cells captured in the scRNA-seq dataset were likely due to valve cell contamination in the ventricular tissue isolations at those earlier stages. We therefore excluded these cells from the lineage analysis. Transferring vascular endothelial cell labels from scRNA-seq data to the spatial data demonstrated that these cells begin to show up at the day 7 stage, which is at the start of coronary vascular development, and are present throughout the myocardium at the day 10 and 14 stages as expected (**Sup. Fig. 2J**). These vascular endothelial cells were also excluded from the lineage analysis since their lineage origin remains debated[29–31].

### Spatiotemporally resolved local cellular heterogeneity in developing cardiac tissue

Ventricular tissue development and morphogenesis in fetal hearts involves spatially patterned regulatory programs. To examine transcriptional differences within the fetal cardiac ventricles, we performed unsupervised clustering of spatial RNA-seq spots and labeled the clusters by anatomical region based on their location in the tissue. Using this analysis, we identified distinct spatial clusters derived from ventricles, atria, valves, and the outflow tract but also distinct layers of ventricular regions including epicardium, compact and trabecular myocardium regions, and endocardium (**Fig. 3A**). Differences in local gene expression can be explained by either difference in cellular composition or cell-type specific gene expression. We used cell type prediction scores for spatial transcriptomes as a proxy for cellular composition and analyzed temporal changes in local cellular composition for the major anatomical regions within ventricular tissue (**Fig. 3B**). We observed a decline of the myocardial progenitor cell population and an increase in the differentiated cardiomyocyte population across stages in all ventricular regions, as expected. The average proportion of endothelial cells decreased in both left and right ventricles, likely due to a reduction in fenestrated trabeculated myocardium layers with developmental time. We observed an increase in the abundance of fibroblast cells with stage in atrial tissue and trabecular ventricular tissue, and a decrease in abundance of fibroblasts in both compact left ventricles and right ventricles. Two to eight cell types contributed to each local transcriptome (number of distinct cell types with a prediction score greater than 5%). We observed a low local cell-type heterogeneity in the valve region across all stages and a high heterogeneity in the trabecular regions at day 7 and day 10 **(Fig. 3C)**. This was likely due to the presence of endocardial cells lining the trabecular myocardium. At day 14, we observed spots with high heterogeneity interspersed in the compact ventricle, which surround vascular bundles **(Fig. 3C, Sup. Fig. 2J)**. Overall, our analysis revealed region specific cell type heterogeneity across the cardiac tissue and supports the idea of collective differentiation and morphogenesis.

**Figure 3:**
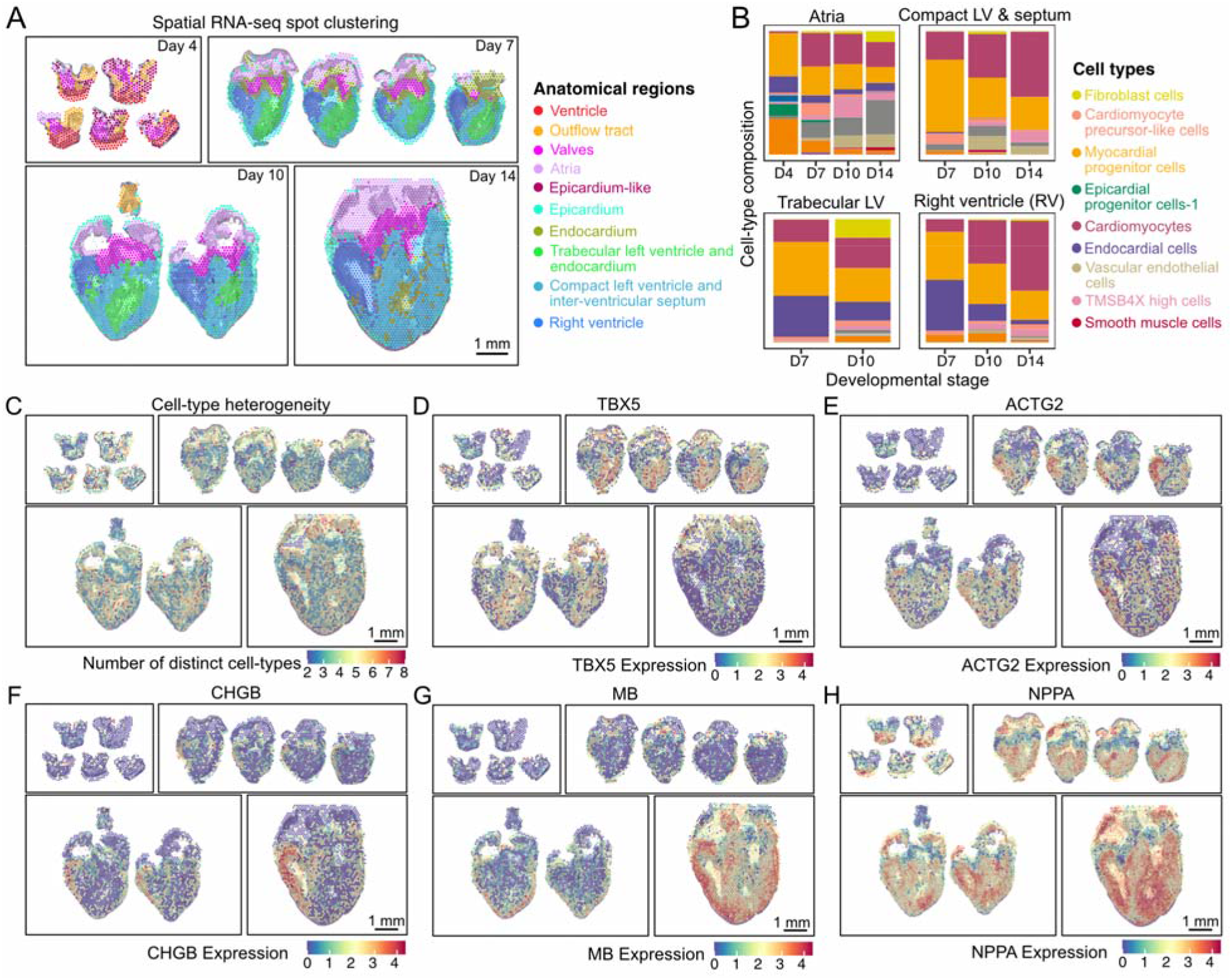
Spatial RNA-seq reveals spatially restricted genes in cardiac tissue during development. **A)** Spatial RNA-seq barcoded spots clustered by gene expression and labeled by tissue anatomical compartment for four developmental stages. **B)** Average cell type composition across various tissue anatomical compartments. **C)** Spatial map showing the cell type heterogeneity for every spot in cardiac tissue across stages. Cell type was estimated by enumerating the number of distinct cell-types with prediction scores greater than 5%. **D-F)** Spatially resolved gene expression for spatially restricted genes differentially expressed between left and right chicken cardiac ventricles. **D)** TBX5 overexpressed in left ventricles on day 7 and day 10. **E)** ACTG2 overexpressed in right ventricles on day 7. **F)** CHGB overexpressed in right ventricles across stages. **G-H)** Spatially resolved normalized gene expression for spatially restricted genes differentially expressed in corresponding ventricular compartments. **G)** MB expressed in compact myocardium on day 10 and day 14. MB expression increases with development stages. **H)** NPPA overexpressed in trabecular myocardium across developmental stages.

To detect region specific markers, we performed differential gene expression analysis between anatomical regions **(Sup. Fig. 4A)**. Interestingly, we found stage dependent transcriptional differences in left and right ventricles at both day 7 and day 10. These differences diminished by day 14 when the main cardiogenesis events were complete. T-box transcription factor 5 (TBX5) expression was mostly restricted to the left ventricle and ACTG2 expression was enriched in the right ventricle on days 7 and 10 (**Fig. 3D&E**). Chromogranin B (CHGB) expression was mostly restricted to the right ventricle from day 7 onwards (**Fig. 3F**). TBX5 specifies the positioning of the left and right ventricular chambers and has been shown to be enriched in the left ventricle of developing chick hearts[32]. We note that the same study by Takeuchi et al. also reported right-ventricle specific expression of TBX20, but our spatial transcriptome data did not corroborate this finding **(Sup. Fig. 4C)**. Unlike spatial RNA-seq data, scRNA-seq datasets from day 7 to 14 did not capture the significant differences in TBX5, ACTG2, and CHGB ventricular expression, although they did seem to trend towards being expressed in a higher proportion of cells in their proposed ventricles **(Sup. Fig. 4B)**. We observed differences in trabeculated versus compact ventricular myocardium at day 7 and day 10 when the transition from trabeculated to compact myocardium is underway. Myoglobin (MB) was spatially restricted to the developing compact myocardial layers across developmental stages (**Fig. 3G**). The emergence of myoglobin in the compact layer of developing myocardium is indicative of cardiomyocytes transitioning to a more mature phenotype with a greater demand for oxygen. By accounting for local cell-type composition, we found that this MB upregulation was a result of both an increase in mature myocytes and increased MB expression in myocyte cells in compact myocardium (**Fig. 3B, Sup. Fig. 4D**). Natriuretic peptide A (NPPA) was spatially restricted to developing trabeculated myocardial layers, as expected because NPPA is a known trabeculated myocardium marker[33] (**Fig. 3H**).

To categorize spatially variable genes (SVGs, genes that correlate with location within a tissue), we used the “markvariogram” method[34] implemented in Seurat-v3 which models spatial transcriptomics data as a mark point process and computes a ‘variogram’ to identify SVGs. This method not only detects genes that are spatially restricted (i.e. gene markers specific to a particular spatially restricted cell type) but also detects ubiquitously expressed genes whose expression correlates with spatial location within anatomical regions. Using this approach, we identified genes spatially restricted to anatomical regions within ventricular tissue (MYH15), atrial tissue (MYH7), and valves (SOX9) across all stages (**Sup. Fig. 4E-G**). SVGs further included BMP4, an epicardial progenitor marker spatially restricted on day 7, fibroblast marker POSTN restricted on days 7 and 10, and vascular endothelial cell marker FABP5 spatially restricted on day 14 (**Sup. Fig. 4H-J**). These SVGs identify functional morphological compartments with distinct transcriptional programs during development.

### A collective cell type and stage dependent role for thymosin beta-4

Unsupervised clustering of single-cell transcriptomes revealed a heterogeneous cell cluster enriched in TMSB4X that contain cells from multiple cardiac lineages and from all four developmental stages (TMSB4X High Cells in **Fig. 1B**, **Sup Fig. 5A**). TMSB4X encodes thymosin beta-4, a well-known secreted small peptide, which plays an important role in actin cytoskeletal organization, cellular motility, survival, and differentiation[35].

Because little is known about the spatiotemporal and cell-type specific expression profile of thymosin beta-4 during cardiogenesis, we investigated the heterogeneity of cellular phenotypes within the TMSB4X cluster in depth. We first collected and re-clustered cells from this cluster and performed differential gene expression analysis to examine cell type composition (**Fig. 4A, Sup. Table 3**). We found that a first subset of these cells (cluster 1) mainly consisted of epicardial progenitor cells expressing TCF21 and endocardial cells expressing endocardial restricted markers IRX6[36] and NPC2[37] (**Fig. 4A,** right). Cluster 1 also contains cardiomyocyte cells expressing MYH15 and ACTC1. A second subset (cluster 2) mainly consisted of vascular smooth muscle-like cells that differentially express ACTA2 (**Fig. 4A**, right). The last subset of these cells (cluster 3) mainly consisted of coronary vascular endothelial-like cells that differentially express FABP5 and TM4SF18 (**Fig. 4A**, right). Interestingly, most of the epicardial progenitors, endocardial cells, and cardiomyocytes in the thymosin beta-4 cluster were detected at early developmental time points, before and during the onset of ventricular compaction (HH24 and HH30), whereas the thymosin beta-4 enriched coronary vascular smooth muscle and vascular endothelial cells were captured at later developmental time points (HH36 and HH40), during the middle and end of ventricular compaction (**Fig. 4A** left). Further analysis revealed a slight increase in thymosin beta-4 expression with developmental stage and highest thymosin beta-4 expression within subcluster 2 (**Sup. Fig. 5B**). One other beta-thymosin, TMSB15B, was expressed but TMSB15B expression did not change with stage **(Sup. Fig. 5D)**.

**Figure 4:**
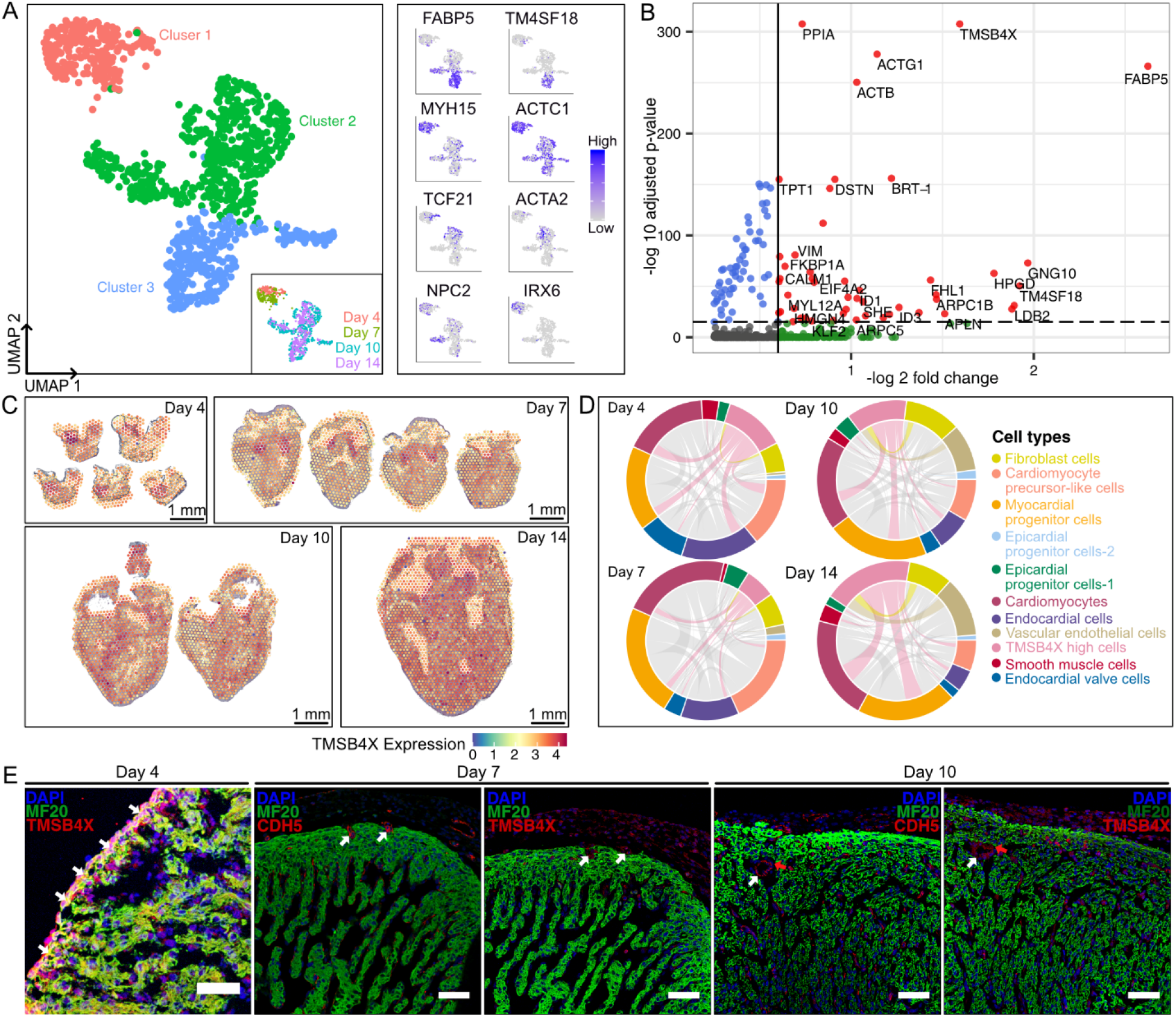
A development stage and cell type dependent role for thymosin beta-4. **A)** UMAP projection of 1075 scRNA-seq TMSB4X high cells clustered by gene expression (left). Inset shows the UMAP projection of TMSB4X high cells labeled by development stage. Feature plots showing expression of gene markers for cell types in the “TMSB4X high cells” cluster: MYH15 and ACTC1 for cardiomyocytes, FABP5 and TM4SF18 for vascular endothelial, ACTA2 for vascular smooth muscle, TCF21 for epicardial lineage, NPC2 and IRX6 for endocardial (right). **B)** Volcano plot showing differentially expressed genes for TMSB4X high scRNA-seq cells. Dotted lines represent thresholds for significantly enriched genes (red): −log_2_ fold change > 0.5 and p-value < 10^−5^. **C)** Spatially resolved thymosin beta-4 (TMSB4X) normalized gene expression across stages in chicken heart spatial RNA-seq data. **D)** Chord diagrams represent cell type proximity maps showing the degree of colocalization of different cell types with TMSB4X high cells within spots in spatial RNA-seq data. **E)** Immunostaining images of chicken heart sections across stages labeled for cardiomyocyte cell marker MF20 (green), endothelial cell marker CDH5 (red), and thymosin beta-4 TMSB4X (red). Representative images of three to four biological replicates. For the day 4 stage, the highest thymosin beta-4 protein expression was observed in the epicardium and the outermost layer of cardiomyocytes (MF20+ and TMSB4X+ colocalization on heart section within the outermost layer of cardiomyocytes). For the day 7 stage, the highest thymosin beta-4 protein expression was detected in developing endothelial-like cells in the myocardium (CDH5+ and TMSB4X+ colocalization on sister sections). For the day 10 stage, the highest thymosin beta-4 protein expression was detected within the vascular bundles of the compact myocardium, which appear to be vascular endothelial cells (CDH5+ and TMSB4X+ colocalization on sister sections within vascular bundles) and vascular smooth muscle cells (CDH5− but TMSB4X+ on sister sections within vascular bundles) (Scale bar = 50μm).

To gain a functional understanding of the TMSB4X high cell cluster, we performed differential gene expression analysis (**Fig. 4B,** Methods). This analysis revealed significant upregulation of genes associated with cytoskeleton organization in the high TMSB4X cell cluster (vimentin (VIM), Rho GTPase (RHOA), actin beta (ACTB), actin gamma 1 (ACTG1), and destin actin depolymerizing factor (DSTN)). In addition, we observed upregulation of calcium binding proteins calmodulin 1 (CALM1) and calmodulin 2 (CALM2), which are associated with cell cycle progression, proliferation, and signaling and have been shown to be activated by thymosin beta-4[38,39]. Our analysis further identified upregulation of peptidylprolyl isomerase A (PPIA) and FK506 binding protein prolyl isomerase 1A (FKBP1A). It is well known that FKBP1A is required for ventricular maturation[40], however, a role for PPIA in cardiogenesis has not previously been reported. Overall, we conclude that TMSB4X enriched cells exhibit a migratory phenotype with increased cytoskeleton organization activity, as confirmed by gene ontology analysis (**Sup. Fig. 5C**).

We used spatially resolved RNA-seq to study TMSB4X expression across the tissue and to characterize the TMSB4X enriched cells in a morphological context. TMSB4X was found to be enriched in atrioventricular valves and the ventricular wall at later stages (**Fig. 4C**). Cell type prediction scores revealed spots comprising cells from the ‘TMSB4X high” cluster in cardiac tissues across stages (**Sup. Fig. 2K**). Spots with a high prediction score for the “TMSB4X high cells” cluster were detected within regions of the outermost layer of myocardium on day 4 and interspersed throughout the compact myocardium at day 14, which most likely represent coronary vascular cells. We next used proximity maps to look for cell type composition of the spots containing the thymosin beta 4 expressing cells and to screen for possible regulatory interactions. As expected, we saw a significant colocalization of thymosin beta enriched cells with cardiomyocytes and endothelial cells in the early stages and vascular-like cells in the later stages (**Fig. 4D, Sup. Fig. 2K**).

To further validate these observations, we performed independent immunostaining experiments (Methods). At the day 4 stage, the highest thymosin beta-4 protein expression was in the epicardium and the outermost layer of cardiomyocytes (MF20+ and TMSB4X+ colocalization) and distributed at lower intensity within the endocardium and trabeculated myocardium. At day 7, the highest thymosin beta-4 protein expression was in the epicardium and developing coronary vascular endothelial cells in the compact myocardial layer (CDH5+ and TMSB4X+ colocalization on sister sections) and at lower intensity in the endocardium but no expression was observed in cardiomyocytes. By day 10, the greatest thymosin beta-4 protein intensity was detected within the vascular bundles of the compact myocardium, which appear to be vascular endothelial cells (CDH5+ and TMSB4X+ colocalization on sister sections within vascular bundles) and vascular smooth muscle cells (CDH5− but TMSB4X+ on sister sections within vascular bundles) as well as lower intensity in the endocardium and epicardium (**Fig. 4E**). Additional immunostaining images split by channel can be found in the supplement (**Sup. Fig. 5E**). Together, our combined spatially resolved transcriptomic and imaging data reveals that TMSB4X plays a ventricular developmental stage and coordinated collective cell-type dependent role across multiple cardiac lineages.

## DISCUSSION

We have combined single-cell and spatial transcriptomics to create a hierarchical map of cellular lineages and their interactions in the developing chicken heart. The dataset spans four development stages, 22,000 single-cell transcriptomes, and 12 spatial gene expression maps. By combining spatial and scRNA-seq assays with novel bioinformatic approaches, we were able to analyze cellular interactions within cardiac tissue, to measure changes in cellular composition with anatomical location, and to quantify anatomically restricted gene expression. Our analysis provides novel insight into several regulatory programs that guide ventricular morphogenesis and the formation of a four chambered fetal heart. Because cellular transitions in complex lineages do not occur in a synchronized manner, the data represents a broad range of cellular states even though we have investigated just four cardiac development stages. We mapped information about lineage differentiation transitions, obtained from pseudotime ordering of single-cell transcriptomes, on spatial maps, and were thereby able to reveal how cellular differentiation and morphological changes co-occur. Our analysis thus indicates that combined spatial and single-cell RNA-sequencing can be used to study both molecular and morphological aspects of development at high spatial and temporal resolution.

The data presented here furthermore provide much needed clarity and insight into the role of thymosin beta 4 in ventricular development, which has been under significant debate in the last 15 years. A pioneering study by Smart et al. demonstrated that conditional knockdowns of thymosin beta-4 in cardiomyocytes give rise to a ventricular non-compaction phenotype in a mouse model, and suggested that cardiomyocytes express thymosin beta-4 in early development in a paracrine manner to induce epicardial progenitor cell migration and ventricular compaction[41]. Our observation of TMSB4X expressing cardiomyocyte and epicardial progenitor cells before the initiation of ventricular compaction is in line with these findings. A more recent study of thymosin beta-4 endothelial-cell specific shRNA knockdowns in fetal mice reported coronary vascular defect phenotypes[42]. In addition, a study of human hearts reported thymosin beta-4 protein expression within vascular endothelial cells as well as epicardium and endocardium but not in cardiomyocytes later in development[43]. Our data is also in line with this prior literature. We observe TMSB4X expression in endocardial and epicardial cells during the beginning of ventricular compaction and in coronary vascular cells during the middle and end of ventricular compaction. Nonetheless, TMSB4X knockout studies have led to discrepancies with prior knockdown studies and have called to question the role of thymosin beta 4 in ventricular development. Whereas Rossdeutsch et al. reported that global thymosin beta 4 knockouts in mice indeed led to a wide range of phenotypes from no abnormalities to lethality[42], another group showed that global, cardiomyocyte-specific, and endothelial specific thymosin beta 4 knockouts in mice were not associated with cardiac abnormalities[44,45]. These discrepancies can potentially be explained by compensatory mechanisms being triggered when thymosin beta-4 is completely ablated in the knockouts, such as the use of other prevalent beta thymosins that have similar functional activity[45–47]. Overall, our study revealed a development stage and cell type dependent role for thymosin beta 4 in which early staged cardiomyocytes, endocardial cells, and epicardial cells express thymosin beta 4 followed by an expression switch to vascular-like cells in the later stages. The small secreted peptide of thymosin beta-4 is therefore capable of coordinating multiple cardiac cell lineages in a developmental spatiotemporal manner and suggests its potential morphogenetic role in directing epicardial and endocardial-derived cell migration into the myocardium to initiate and maintain ventricular compaction and maturation. To the best of our knowledge, no studies to date have revealed this switch in dominant thymosin beta-4 expression across ventricular development stage and cell type. Beta thymosins have regenerative potential as pleiotropic factors in adult myocardial infarction models[35,48,49], and therefore a deeper understanding of how beta thymosins influence cardiac cell types during ventricular development could lead to novel treatments of myocardial infarctions and congenital cardiac diseases.

Our analysis demonstrates several ways in which spatial RNA-seq can be used to improve the robustness of scRNA-seq, and vice versa, how scRNA-seq can be used to improve the robustness of spatial RNA-seq. scRNA-seq requires surgical isolation of tissues of interest, which is often experimentally challenging, making scRNA-seq prone to contamination by cells from adjacent tissues. Matched spatial RNA-seq can be used to identify such cellular contamination. For example, using spatial RNA-seq we were able to identify contamination of valve endocardial cells in ventricular scRNA-seq data collected at day 4 and day 7 stages. In addition to cellular contamination, scRNA-seq is sensitive to contamination by cell free RNA released from cells that lyse during tissue dissociation[50,51]. We found that spatial RNA-seq is helpful to identify sources of cell-free RNA contamination, e.g. contamination by hemoglobin RNA from red blood cells and troponin RNA from cardiomyocytes (data not shown). Spatial RNA-seq conversely suffers from limitations due to local ensemble averaging that can be mitigated with matched scRNA-seq data. Spot tissue expression profiles obtained by spatial RNA-seq are a weighted mix of the single-cell expression profiles of the cells in the spot. ScRNA-seq data for matched tissue can be used to obtain the weights of scRNA-seq derived cell types, thereby effectively increasing the resolution of spatial RNA-seq. This analysis enables a measurement of the local cellular composition that is more accurate than a cell type composition obtained by scRNA-seq alone because it does not require tissue dissociation which can introduce cell-type biases.

We note that previous scRNA-seq studies have studied the development of human and mouse hearts[52–54]. These studies implemented low-throughput scRNA-seq methodologies and consequently suffered from limitations in cell type resolution. A very recent study combined spatial transcriptomics combined with scRNA-seq to construct a spatiotemporal organ-wide gene expression and cell atlas of developing human hearts[55] at three developmental stages in the first trimester: 4.5–5, 6.5, and 9 post-conception weeks (PCW). Although this spatial gene expression atlas from human hearts was resolved to the single-cell level using scRNA-seq data, the study has limitations with respect to detecting rare cell types and identifying spatiotemporal cell-cell interactions due to the limited cell types being detected (total of only 3,717 cells), limited genes being used in the In-Situ Sequencing (ISS) panel, and low resolution of the spatial transcriptomic technique (3,115 spots containing ~30 cells per spot). In comparison to chick heart staging, the human heart stages of this study were the following: *i)* 4.5-5pcw ≈ HH28-31, *ii)* 6.5pcw ≈ HH34-37, and *iii)* 9pcw ≈ HH40-later[3,56]. Therefore, our study was able to probe earlier stages of development (HH21-24). This early chamber formation stage consists of major morphogenetic events including the initiation of ventricular septation and corresponding four-chambered heart structure[3].

In conclusion, we have combined single-cell and spatial transcriptomics to explore the early development of chicken hearts at high molecular, spatial and temporal resolution. We constructed a single cell, spatially resolved gene expression atlas, uncovered several novel regulatory mechanisms, and demonstrate that spatiotemporal single-cell RNA sequencing can be used to study the interplay between cellular differentiation and morphogenesis.

## METHODS

### Sample preparation for single-cell transcriptomics

Fertile bovans brown chicken (Gallus *gallus*) eggs were incubated in a standard humidity and temperature-regulated egg incubator until the embryonic day of interest. Ventricles were isolated aseptically, placed in ice cold Hank’s Balanced Salt Solution, and minced into 1-2mm pieces. Six dozen day 4 (HH21-24) whole ventricles, four dozen day 7 (HH30-31) left and right ventricles, three dozen day 10 (HH35-36) left and right ventricles, and one dozen day 14 (HH40) left and right ventricles were pooled respectively for a total of seven samples to be analyzed via single-cell RNA sequencing. Day 4 (HH21-24) and day 7 (HH30-31) ventricular samples were digested in 1.5mg/mL collagenase type II/ dispase (Roche) for one cycle of 20 minutes and one cycle of 10 minutes under mild agitation at 37°C. Day 10 (HH35-36) and day 14 (HH40) ventricular samples were digested in 300U/mL collagenase type II (Worthington Biochemical Corporation) for four cycles of 10 minutes under mild agitation at 37°C. At the end of the digestions, the cells were passed through a 40μm filter and centrifuged into a pellet. To remove most blood contaminants, samples were resuspended in an ammonium-chloride-potassium red blood cell lysis buffer (Thermo Fisher Scientific) and centrifuged again. Samples were then resuspended in phosphate buffered saline with 0.04% bovine serum albumin (Thermo Fisher Scientific) at 1⎕×⎕10^6^ cells per ml.

### Single-cell RNA sequencing library preparation

4,000-5,000 cells per sample (for day 4 and day 7 samples) and 2,000-3,000 cells per sample (for day 10 and day 14 samples) were targeted on the Chromium platform (10X Genomics) using one lane per time point. Single-cell mRNA libraries were built using the Chromium Next GEM Single Cell 3’ Library Construction V3 Kit for day 4 and day 7 samples and Chromium Next GEM Single Cell 3’ Library Construction V2 Kit for day 10 and day 14 samples, libraries sequenced on an Illumina NextSeq 500/550 using 75 cycle high output kits (Index 1⎕= 8, Read 1⎕=⎕28, and Read 2⎕=⎕55) for day 4 and day 7, 75 cycle high output kits (Index 1⎕= 8, Read 1⎕=⎕26, and Read 2⎕=⎕98) for day 10, and 75 cycle high output kits (Index 1⎕= 8, Read 1⎕=U26, and Read 2⎕=⎕58) for day 14. Sequencing data was aligned to chicken reference (assembly: GRCg6a) using the Cell Ranger 3.0.2 pipeline (10X Genomics).

### Reference genome and annotation

*Gallus gallus* genome and gene annotations (assembly: GRCg6a) were downloaded from Ensembl genome browser 97 and processed using the Cell Ranger 3.0.2 (10X Genomics) pipeline’s mkref command. The reference was then used in the Cell Ranger “count” command to generate expression matrices.

### scRNA-seq data processing, batch correction, clustering, cell-type labeling, and data visualization

Cells with less than 1200 UMIs or 200 unique genes, or more than 30 percent mitochondrial transcripts were excluded from the dataset. The remaining 22315 cells from 7 samples were transformed, normalized, and scaled using the Seurat V3 package, and then used for batch correction. Scanorama[4] was used for dataset integration and batch correction using transformed and normalized expression values. We used the batch corrected values for further processing and analysis. Seurat-v3 package was used to select top variable genes for scRNA-seq clustering. We used the FindVariableFeatures function in Seurat to choose the top 2000 highly variable genes from our dataset using the “vst” selection method. We then performed mean centering and scaling, followed by principal component analysis (PCA) on a matrix composed of cells and batch-corrected scanorama-output values, and reduced the dimensions of our data to the top 20 principal components. Uniform Manifold Approximation and Projection (UMAP) was initialized in this PCA space to visualize the data on reduced UMAP dimensions. The cells were clustered on PCA space using the Shared Nearest Neighbor (SNN) algorithm implemented as FindNeighbors and FindClusters in Seurat-v3. The method returned 17 cell clusters which were then visualized on UMAP space using the DimPlot command as shown in **Figure 1**. Cell-type specific canonical gene markers were used to label clusters differentially expressing those markers. To accurately label individual clusters, Wilcox test was performed to find differentially expressed genes for each cluster. We used the FindAllMarkers function in Seurat to get a list of differentially expressed genes for each cluster. Gene expression was visualized using FeaturePlot, DoHeatMap, and VlnPlot functions from Seurat-v3. Cells were grouped into lineages using gene markers and then used for trajectory construction and pseudotime analysis.

### Sample preparation for 10X Visium spatial transcriptomics platform

Whole hearts were isolated using aseptic technique and placed in ice cold sterile Hank’s Balanced Salt Solution and then blood was carefully removed by perfusing the hearts through the apex. Fresh tissues were immediately embedded in Optimal Cutting Compound (OCT) media and frozen in liquid-nitrogen-cooled isopentane bath, cut into 10μm sections using Thermo Scientific CryoStar NX50 cryostat, and mounted on 10X Visium slides, which were pre-cooled to −20°C.

### 10X Visium spatial transcriptomics library preparation

Tissue sections from fresh frozen chicken embryonic hearts were mounted for 4 stages (five sections from day 4- HH24, four sections for day 7- HH31, two sections for day 10- HH36, and one section for day 14- HH40) with one stage per capture area on a 10x Visium gene expression slide containing four capture areas with 5000 barcoded RNA spots printed per capture area. The sample for day 4 had three biological replicates with two of them having technical replicates and the samples for day 7 and day 10 had four and two biological replicates, respectively. Spatially tagged cDNA libraries were built using the 10x Genomics Visium Spatial Gene Expression 3’ Library Construction V1 Kit. Optimal permeabilization time for 10μm thick chicken heart sections was found to be 12 minutes using 10X Genomics Visium Tissue Optimization Kit. H&E stained heart tissue sections were imaged using Zeiss PALM MicroBeam laser capture microdissection system and the images were stitched and processed using Fiji ImageJ software. cDNA libraries were sequenced on an Illumina NextSeq 500/550 using 150 cycle high output kits (Read 1⎕= 28, Read 2⎕=⎕120, Index 1⎕=⎕10, and Index 2⎕=⎕10). Fluidigm frames around the capture area on the Visium slide were aligned manually and spots covering the tissue were selected using Loop Browser 4.0.0 software (10X Genomics). Sequencing data was then aligned to the chicken reference genome using the Space Ranger 1.0.0 pipeline to derive a feature spot-barcode expression matrix (10X Genomics).

### Spatial RNA-seq data processing, integration, and visualization

Over 6,800 barcoded spatial spots from four 10X Visium capture areas were transformed and normalized using the Seurat V3.2 package, and then used for batch correction. Seurat-v3.2 package was used to select top variable genes for spatial RNA-seq clustering. We used the FindVariableFeatures function in Seurat to choose the top 2000 highly variable genes from our dataset using the “vst” selection method. We then performed mean centering and scaling, followed by principal component analysis (PCA) on a matrix composed of spots and gene expression (UMI) counts, and reduced the dimensions of our data to the top 20 principal components. Uniform Manifold Approximation and Projection (UMAP) was initialized in this PCA space to visualize the data on reduced UMAP dimensions. The spots were clustered on PCA space using the Shared Nearest Neighbor (SNN) algorithm implemented as FindNeighbors and FindClusters in Seurat v3.2 with parameters k = 30, and resolution = 0.5. The method returned spot clusters representing anatomical regions in the tissues, which were then visualized on UMAP space using the SpatialDimPlot command as shown in **Figure 1**. To accurately label anatomical regions, Wilcox test was performed to find differentially expressed genes for each region. We used the FindAllMarkers function in Seurat with its default parameters to get a list of differentially expressed genes for each cluster. Gene expression was visualized using SpatialFeaturePlot function from Seurat v3.2. An anchor-based integration method implemented in Seurat-v3.2 was used for integration of spatial RNA-seq data with time matched scRNA-seq data using FindIntegrationAnchors command and then cell type labels were transferred to spatial data using TransferData command. Cell type prediction values for Spatial RNA-seq spots were saved as an assay and used for further analysis. Cell type colocalization values were calculated by counting cell type pair abundances in spatial RNA-seq spots. Only cell types with top four prediction scores in each spot were included. We also constructed a cell-spot similarity map by transferring cell barcode IDs to spatial barcoded spots. The cell-spot similarity matrix containing scRNA-seq cell similarity prediction for each spot in scRNA-seq data, which we further used to estimate pseudotime for spatial RNA-seq spots.

### Pseudotime analysis and trajectory construction

We used PHATE[7] (Potential of Heat-diffusion for Affinity-based Transition Embedding) to visualize developmental trajectories because of its ability to learn and maintain local and global distances in low dimensional space. We reclustered cells from individual lineages, performed PHATE dimension reduction on scanorama integrated values, and used PHATE1 dimension as a proxy for development time. PHATE reduction was performed using the phate command implemented in the R package: phateR. We also used monocle-v2[20,21] to order the cells in epicardial, endocardial, and myocardial lineages along pseudotime and reconstruct lineage trajectories. We filtered the genes detected in our dataset and retained the top 2,000 highly variable genes calculated using Seurat-v3 in our monocle analysis. We further filtered these genes to genes differentially expressed in cell type subclusters within the lineage using differentialGeneTest command and then reduced the dimension of our data using the DDRTree method. We used the ReduceDimension function in monocle-v2 to reduce the dimension to two DDRTree components, which is then used to define a pseudotime scale. The cells were then ordered along pseudotime using monocle’s orderCells command, and the root of the trees was defined as the branch with maximum cells from Day 4 samples. The gene expression changes along pseudotime based trees were then visualized using PseudotimeHeatMap command in monocle-v2. To estimate pseudotime for spatial maps, we used the spot-cell similarity matrix and estimated pseudotime for individual lineages in spatial RNA-seq data. We defined spot pseudotime for a lineage as the average of scRNA-seq pseudotime (PHATE1 dimension or monocle pseudotime) for cells having a non-zero similarity prediction with that spot. This spot pseudotime was then visualized on spatial maps using SpatialFeaturePlot command in Seurat-v3.2.

### Enrichment analysis for Gene Ontology (GO)

We used the genes differentially expressed between clusters in individual lineages to perform enrichment analysis for Gene Ontology. We selected significant genes using their p-value scores from differential expression test and used the classic fisher test implemented in topGO[57] R package for enrichment analysis of GO terms representing various Biological Processes (BP).

### Immunohistochemistry Assays

Whole hearts were isolated using aseptic technique and placed in ice cold sterile Hank’s Balanced Salt Solution and then blood was carefully removed by perfusing the hearts through the apex. For day 7 (HH30), day 10 (HH36), and day 14 (HH40) heart samples after perfusion, tissues were fixed in 4% paraformaldehyde for about 16-24 hours at 4°C. Samples were then processed, embedded in paraffin, cut into 6μm sections using a microtome, and mounted onto histological glass slides. Slides were incubated for twenty minutes at 58°C to melt paraffin, washed three times in xylene for three minutes each, and then placed in decreasing ethanol concentrations to rehydrate slides. Samples then underwent an antigen retrieval step via incubation in 1X citrate buffer for at least 10 minutes at 95°C. Samples were then permeabilized in 0.3% Triton X-100 in tris buffered saline for fifteen minutes, washed three times in 0.05% Tween-20 in tris buffered saline (TBST), blocked for one hour at room temperature in blocking buffer (1% bovine serum albumin and 5% goat serum in TBST), washed once in TBST, and incubated in primary antibodies diluted in antibody solution (1% bovine serum albumin in TBST) overnight at 4°C. Primary antibodies used were mouse anti-MF20 antibody (1:100, 14650382, Invitrogen), rabbit anti-TMSB4X antibody (1:200, ab14335, Abcam), and rabbit anti-CDH5 antibody (1:200, ab33168 Abcam). After overnight primary incubation, samples were washed three times in TBST and then incubated in secondary antibodies for one hour at room temperature. The secondary antibodies were goat anti-mouse 488 (1:500, A10684, Invitrogen) and goat anti-rabbit 555 (1:500, A21430, Invitrogen). Lastly, samples were washed in TBST, stained with DAPI (1:1000, Thermofisher), and mounted. Images were acquired using a Zeiss 880 inverted confocal microscope.

For day 4 (HH24) heart samples after perfusion, fresh tissues were immediately embedded in Optimal Cutting Compound (OCT) media and frozen in liquid nitrogen cooled isopentane, cut into 6μm sections using a Thermo Scientific CryoStar NX50 cryostat, and mounted on −20°C cooled histological glass slides. The mounted day 4 (HH24) cryosections were thawed at 37°C to melt the OCT and immediately placed in 4% paraformaldehyde for 20 minutes at room temperature. Samples were then washed three times in phosphate buffered saline (DPBS), permeabilized with 0.3% Triton X-100 in DPBS for fifteen minutes, washed three times in DPBS, and then blocked in blocking buffer (1% bovine serum albumin and 5% goat serum in DPBS) for one hour at room temperature. Following the blocking step, samples were washed in DPBS, and then incubated overnight at 4°C in primary antibodies diluted in antibody solution (1% bovine serum albumin in DPBS). Primary antibodies were mouse anti-MF20 antibody (1:100, 14650382, Invitrogen) and rabbit anti-TMSB4X antibody (1:200, ab14335, Abcam). After overnight primary incubation, samples were washed three times in DPBS and then incubated in secondary antibodies for one hour at room temperature. The secondary antibodies were goat anti-mouse 488 (1:500, A10684, Invitrogen) and goat anti-rabbit 555 (1:500, A21430, Invitrogen). Lastly, samples were washed, stained with DAPI (1:1000, Thermofisher), and mounted. Images were acquired using a Zeiss 880 inverted confocal microscope.

## ACKNOWLEDGEMENTS

We would like to thank Peter Schweitzer and the Cornell Genomics Center for help with single-cell and spatial sequencing assays, the Cornell Bioinformatics facility for assistance with bioinformatics, and the Cornell Imaging facility for assistance with imaging assays. We thank the members of the Butcher and De Vlaminck labs for valuable discussion. This work was supported by R01HL143247 (to J.B. and I.D.V), R33CA235302 (to I.D.V.), R21AI133331 (to I.D.V.), and DP2AI138242 (to I.D.V.).

## DATA AVAILABILITY

The sequencing data discussed in this publication have been deposited in NCBI’s Gene Expression Omnibus[58] and are accessible through GEO Series accession number GSE149457 (https://www.ncbi.nlm.nih.gov/geo/query/acc.cgi?acc=GSE149457). H&E stained tissue images matched to spatial RNA-seq datasets and scripts have been made available on GitHub (https://github.com/madhavmantri/chicken_heart).

## REFERENCES

1. Buckingham M, Meilhac S, Zaffran S. Building the mammalian heart from two sources of myocardial cells. Nat Rev Genet 2005; 6:826–835.

2. Dunwoodie SL. Combinatorial signaling in the heart orchestrates cardiac induction, lineage specification and chamber formation. Semin Cell Dev Biol 2007; 18:54–66.

3. Martinsen BJ. Reference guide to the stages of chick heart embryology. Dev Dyn 2005; 233:1217–1237.

4. Hie B, Bryson B, Berger B. Efficient integration of heterogeneous single-cell transcriptomes using Scanorama. Nat Biotechnol 2019.

5. Stuart T, Butler A, Hoffman P, et al. Comprehensive integration of single cell data. 2018:1–24.

6. Butler A, Hoffman P, Smibert P, Papalexi E, Satija R. Integrating single-cell transcriptomic data across different conditions, technologies, and species. Nat Biotechnol 2018; 36:411–420.

7. Moon KR, van Dijk D, Wang Z, et al. Visualizing structure and transitions in high-dimensional biological data. Nat Biotechnol 2019.

8. Wu SP, Dong XR, Regan JN, Su C, Majesky MW. Tbx18 regulates development of the epicardium and coronary vessels. Dev Biol 2013; 383:307–320.

9. Xavier-Neto J, Sousa Costa ÂM, Figueira ACM, et al. Signaling through retinoic acid receptors in cardiac development: Doing the right things at the right times. Biochim Biophys Acta - Gene Regul Mech 2015; 1849:94–111.

10. Guadix JA, Ruiz-Villalba A, Lettice L, et al. Wt1 controls retinoic acid signalling in embryonic epicardium through transcriptional activation of Raldh2s. Development 2011; 138:1093–1097.

11. Kadomatsu K, Bencsik P, Görbe A, et al. Therapeutic potential of midkine in cardiovascular disease. Br J Pharmacol 2014; 171:936–944.

12. Kadomatsu K,…SK-TJ of, 2013 undefined. The heparin-binding growth factor midkine: the biological activities and candidate receptors. academic.oup.com.

13. Wang B, Yang W, McKittrick J, Meyers MA. Keratin: Structure, mechanical properties, occurrence in biological organisms, and efforts at bioinspiration. Prog Mater Sci 2016; 76:229–318.

14. Velecela V, Torres-Cano A, Garcıá-Melero A, et al. Epicardial cell shape and maturation are regulated by Wt1 via transcriptional control of Bmp4. Dev 2019; 146.

15. Dupuis LE, Kern CB. Small leucine-rich proteoglycans exhibit unique spatiotemporal expression profiles during cardiac valve development. Dev Dyn 2014; 243:601–611.

16. Tidball JG. Distribution of collagens and fibronectin in the subepicardium during avian cardiac development. Anat Embryol (Berl) 1992; 185:155–162.

17. Benesh EC, Miller PM, Pfaltzgraff ER, et al. Bves and NDRG4 regulate Directional epicardial cell migration through autocrine extracellular matrix deposition. Mol Biol Cell 2013; 24:3496–3510.

18. Snider P, Standley KN, Wang J, Azhar M, Doetschman T, Conway SJ. Origin of cardiac fibroblasts and the role of periostin. Circ Res 2009; 105:934–47.

19. Bassat E, Mutlak YE, Genzelinakh A, et al. The extracellular matrix protein agrin promotes heart regeneration in mice. Nature 2017; 547:179–184.

20. Trapnell C, Cacchiarelli D, Grimsby J, et al. The dynamics and regulators of cell fate decisions are revealed by pseudotemporal ordering of single cells. Nat Biotechnol 2014; 32:381–386.

21. Qiu X, Mao Q, Tang Y, et al. Reversed graph embedding resolves complex single-cell trajectories. Nat Methods 2017; 14:979–982.

22. Horrillo A, Porras G, Ayuso MS, González-Manchón C. Loss of endothelial barrier integrity in mice with conditional ablation of podocalyxin (Podxl) in endothelial cells. Eur J Cell Biol 2016; 95:265–276.

23. Privratsky JR, Newman PJ. PECAM-1: Regulator of endothelial junctional integrity. Cell Tissue Res 2014; 355:607–619.

24. Lee NY, Blobe GC. The interaction of endoglin with beta -arrestin2 regulates transforming growth factor-beta -mediated ERK activation and migration in endothelial cells. J Biol Chem 2007.

25. Terai Y, Abe M, Miyamoto K, et al. Vascular smooth muscle cell growth-promoting factor/F-spondin inhibits angiogenesis via the blockade of integrin◻?v?3 on vascular endothelial cells. J Cell Physiol 2001; 188:394–402.

26. Akazawa H, Komuro I. Cardiac transcription factor Csx/Nkx2-5: Its role in cardiac development and diseases. Pharmacol Ther 2005; 107:252–268.

27. Franco D, Campione M. The role of Pitx2 during cardiac development: Linking left-right signaling and congenital heart diseases. Trends Cardiovasc Med 2003; 13:157–163.

28. Nelson DO, Lalit PA, Biermann M, et al. Irx4 Marks a Multipotent, Ventricular-Specific Progenitor Cell. Stem Cells 2016; 34:2875–2888.

29. Wu B, Zhang Z, Lui W, et al. Endocardial cells form the coronary arteries by angiogenesis through myocardial-endocardial VEGF signaling. Cell 2012; 151:1083–1096.

30. Perez-Pomares JM, Carmona R, González-Iriarte M, Atencia G, Wessels A, Muñoz-Chapuli R. Origin of coronary endothelial cells from epicardial mesothelium in avian embryos. Int J Dev Biol 2002; 46:1005–1013.

31. Katz TC, Singh MK, Degenhardt K, et al. Distinct Compartments of the Proepicardial Organ Give Rise to Coronary Vascular Endothelial Cells. Dev Cell 2012; 22:639–650.

32. Takeuchi JK, Ohgi M, Koshiba-Takeuchi K, et al. Tbx5 specifies the left/right ventricles and ventricular septum position during cardiogenesis. Development 2003; 130:5953–5964.

33. Jensen B, van der Wal AC, Moorman AFM, Christoffels VM. Excessive trabeculations in noncompaction do not have the embryonic identity. Int J Cardiol 2017; 227:325–330.

34. Edsgärd D, Johnsson P, Sandberg R. Identification of spatial expression trends in single-cell gene expression data. Nat Methods 2018.

35. Bock-Marquette I, Saxena A, White MD, DiMaio JM, Srivastava D. Thymosin β4 activates integrin-linked kinase and promotes cardiac cell migration, survival and cardiac repair. Nature 2004; 432:466–472.

36. Mummenhoff J, Houweling AC, Peters T, Christoffels VM, Rüther U. Expression of Irx6 during mouse morphogenesis. Mech Dev 2001; 103:193–195.

37. Anderson C, Hill B, Lu HC, et al. A 3D molecular atlas of the chick embryonic heart. Dev Biol 2019; 456:40–46.

38. Tsang WY, Spektor A, Luciano DJ, et al. CP110 cooperates with two calcium-binding proteins to regulate cytokinesis and genome stability. Mol Biol Cell 2006; 17:3423–3434.

39. Galoyan AA, Gurvitis BYa, Shuvalova LA, Davis MT, Shively JE, Lee TD. A hypothalamic activator of calmodulin-dependent enzymes is thymosin beta 4 (1-39). Neurochem Res 1992; 17:773–7.

40. Chen H, Zhang W, Sun X, et al. Fkbp1a controls ventricular myocardium trabeculation and compaction by regulating endocardial Notch1 activity. Dev 2013; 140:1946–1957.

41. Smart N, Risebro CA, Melville AAD, et al. Thymosin β4 induces adult epicardial progenitor mobilization and neovascularization. Nature 2007; 445:177–182.

42. Rossdeutsch A, Smart N, Dubé KN, Turner M, Riley PR. Essential Role for thymosin β4 in regulating vascular smooth muscle cell development and vessel wall stability. Circ Res 2012; 111.

43. Saunders V, Dewing JM, Sanchez-Elsner T, Wilson DI. Expression and localisation of thymosin beta-4 in the developing human early fetal heart. PLoS One 2018; 13:1–11.

44. Banerjee I, Zhang J, Moore-Morris T, et al. Thymosin beta 4 is dispensable for murine cardiac development and function. Circ Res 2012; 110:456–464.

45. Sribenja S, Wongkham S, Wongkham C, Yao Q, Chen C. Roles and mechanisms of β-thymosins in cell migration and cancer metastasis: An update. Cancer Invest 2013; 31:103–110.

46. Huff T, Müller CSG, Otto AM, Netzker R, Hannappel E. β-thymosins, small acidic peptides with multiple functions. Int J Biochem Cell Biol 2001; 33:205–220.

47. Dhaese S, Vandepoele K, Waterschoot D, et al. The Mouse Thymosin Beta15 Gene Family Displays Unique Complexity and Encodes A Functional Thymosin Repeat. J Mol Biol 2009; 387:809–825.

48. Hinkel R, El-Aouni C, Olson T, et al. Thymosin beta4 is an essential paracrine factor of…[Circulation. 2008] - PubMed result. Circulation 2008; 117:2232–40.

49. Hinkel R, Bock-Marquette I, Hazopoulos AK, Kupatt C. Thymosin β4: a key factor for protective effects of eEPCs in acute and chronic ischemia. Ann N Y Acad Sci 2010; 1194:105–111.

50. Fleming SJ, Marioni JC, Babadi M. CellBender remove-background: a deep generative model for unsupervised removal of background noise from scRNA-seq datasets. biorxiv.org 2019.

51. Young MD, Behjati S. SoupX removes ambient RNA contamination from droplet based single cell RNA sequenc-ing data. biorxiv.org 2018.

52. DeLaughter D, Bick A, Wakimoto H, cell DM-D, 2016 undefined. Single-cell resolution of temporal gene expression during heart development. Elsevier.

53. Li G, Xu A, Research SW-C, 2017 undefined. Transcriptomic Profiling Maps Anatomically Patterned Subpopulations Among Single Embryonic Cardiac Cells. Am Hear Assoc.

54. Cui Y, Zheng Y, Liu X, et al. Single-Cell Transcriptome Analysis Maps the Developmental Track of the Human Heart. Cell Rep 2019; 26:1934–1950.e5.

55. Asp M, Giacomello S, Larsson L, et al. A Spatiotemporal Organ-Wide Gene Expression and Cell Atlas of the Developing Human Heart. Cell 2019.

56. Lindsey SE, Butcher JT, Yalcin HC. Mechanical regulation of cardiac development. Front Physiol 2014; 5 AUG.

57. Alexa, A; Rahnenfuhrer J. topGO: Enrichment Analysis for Gene Ontology. R package version 2.37.0. Rahnenfuhrer 2019.

58. Edgar R. Gene Expression Omnibus: NCBI gene expression and hybridization array data repository. Nucleic Acids Res 2002.

