## Supplemental Materials for "Spatiotemporal single-cell RNA sequencing of developing hearts reveals interplay between cellular differentiation and morphogenesis"

**Supplemental Table 1: Canonical gene markers for scRNAseq cell types (Section 1)**

| **Gene name (Symbol)** | **Cell type(s)** | **Details and references** |
| --- | --- | --- |
| Hemoglobin subunit zeta (HBZ) | Erythrocytes | Present in embryonic staged erythrocytes (Brittain et al 2002, Molecular Aspects of Medicine) |
| Colony stimulating factor 1 receptor (CSF1R) | Macrophages | Macrophage/monocyte specific marker (Jones et al 2013, Organogenesis) |
| Interferon alpha- inducible protein 6 (IFI6) | Dendritic cells | Dendritic cell specific marker (Chan et al 2006, Nature Medicine) |
| SRY-box 9 (SOX9) | Valve endocardial cells | Important transcription factor implicated in valve development (Garside et al 2015, Development) |
| Transmembrane 4 L six family member 18 (TM4SF18) | Vascular endothelial cells | Tetraspanin shown to be an important modulator of angiogenesis. (Page et al 2019, Cell Reports and Margadant 2020, Angiogenesis) |
| Fatty acid binding protein-5 (FABP5) | Vascular endothelial cells | FABP5 plays a role in vascular endothelial migration, proliferation, and overall angiogenesis (Wei Yu et al 2016, Angiogenesis) |
| Vascular endothelial cadherin (CDH5) | Vascular endothelial cells, endocardial cells | Well-known endothelial marker (Dejana et al 2004, Nature Reviews Molecular Cell Biology) |
| NPC Intracellular Cholesterol Transporter 2 (NPC2) | Endocardial cells | An established endocardium-specific marker found in developing chick hearts (Anderson et al 2019, Developmental Biology) |
| Transcription factor 21 (TCF21) | Epicardial progenitor cells-1 and 2 | Required for epicardial specification and maturation (Tandon et al 2013, Development) |
| Agrin (AGRN) | Epicardial progenitor cells 2, endocardial valve cells, endocardial cells, vascular endothelial cells | Glycoprotein shown to be expressed by cardiac endothelial cells during development that is implicated in inducing cardiomyocyte proliferation and dedifferentiation (Bassat et al 2017, Nature) |
| Aldehyde dehydrogenase 1 family member A2 (ALDH1A2) | Epicardial progenitor  cells-1 | An enzyme that facilitates the production of retinoic acid (RA). RA signaling is vital to ventricular development and helps maintain epicardial cells in an undifferentiated state to delay smooth muscle differentiation prior to coronary endothelial network formation (Xavier-Neto et al 2015, Biochimca et Biophysica Acta). This enzyme is regulated by Wilm’s Tumor 1 (WT1), a well-known epicardium regulating transcription factor (Antonio Guadix et al 2011, Development) |
| Bone morphogenetic protein 4 (BMP4) | Epicardial progenitor  cells-1 | A key regulator of epicardial maturation. Wt1 knockout mice have an upregulated BMP4 signaling which prevents the proepicardium from becoming a mature phenotype. Therefore, when present, maintains an epicardial progenitor-like phenotype (Velecela et al 2019, Development) |
| Decorin (DCN) | Fibroblast cells, epicardial progenitor cells-1, epicardial progenitor cells-2, vascular smooth muscle cells | Proteoglycan that binds to collagen fibers and is an important component of structural extracellular matrix production (Weber et al 1996, J. of Biol. Chem.) |
| Midkine (MDK) | Epicardial progenitor cells-1 | Secreted growth factor that responds to retinoic acid signaling. Well-known to promote cell migration and also angiogenesis (Kadomatsu et al 2014 Br J Pharmacol) and (Kadomatsu et al 2013, J of BIochemistry) |
| Collagen type 1 alpha 1 chain (COL1A1) | Fibroblast cells | Fibrillar structural extracellular matrix protein. Produced by cardiac fibroblast cells (Pan et al 2013, Plos One) |
| Smooth muscle actin (ACTA2) | Vascular smooth muscle cells | Well-known smooth muscle cell marker (Yuan et al 2015, Braz J of Cardiovascular Surgery) |
| Iroquois Homeobox 4 (IRX4) | Myocardial progenitor cells, cardiomyocytes | Transcription factor implicated in ventricular specific myocardial progenitor differentiation (Nelson et al 2016, Stem Cells) |
| NK2 homeobox 5 (NKX2-5) | Myocardial progenitor cells, cardiomyocytes | Important transcription factor implicated in myocardial progenitor differentiation (Akazawa et al 2005, Pharmacology and Therapeutics) |
| Myosin light chain 10 (MYL10) | Cardiomyocytes | Paralog of MYL2 which is a ventricular specific myosin light chain. |
| Cardiac type troponin T2 (TNNT2) | Myocardial progenitor cells, Cardiomyocytes | Cardiomyocyte-specific troponin |
| Thymosin beta -4 (TMSB4X) | TMSB4X high cells | G-actin monomer sequestering protein, which plays an important role in actin cytoskeletal organization, cellular motility, survival, and differentiation (Bock-Marquette et al Nature 2004). |

**Supplemental Table 2: Additional gene markers for trajectory analysis (Section 2)**

| **Gene name (Symbol)** | **Cell type(s)** | **Details and references** |
| --- | --- | --- |
| **Endocardial Lineage** | | |
| Podocalyxin (PODXL) | Endocardial cell subclusters C1-C3 | Endothelial specific mucin that facilitates endothelial barrier and integrity (Horrillo et al 2016, European Journal of Cell Biology) |
| Platelet and endothelial cell adhesion molecule 1 (PECAM1) | Endocardial cell subclusters C1-C3 | Well-known endothelial specific marker. Modulates the integrity of endothelial barriers (Privratsky et al 2014, Cell and Tissue Research) |
| Retinoic acid receptor responder 1 (RARRES1) | Endocardial cell subclusters C1-C3 | Becomes upregulated in the presence of retinoic acid receptors and therefore involved in retinoic acid signaling |
| Endoglin (ENG) | Endocardial cell subclusters C1-C3 | Vascular endothelial glycoprotein that facilitates angiogenesis and involved in transforming growth factor beta signaling (Lee et al 2007, Journal of Biological Chemistry) |
| **Epicardial Lineage** | | |
| Lumican (LUM) | Epicardial cell subcluster C2 | Small leucine rich proteoglycan shown to be expressed in the outermost epithelial-like layer of fetal epicardium (Dupuis et al 2013, Developmental Dynamics). |
| Keratin 7 (KRT7) | Epicardial cell subclusters C1 and C2 | Fibrous filament structural proteins found in outermost epithelial-like layers of various organs (Wang et al 2016, Progress in Material Science). |
| T-box transcription factor 18 (TBX18) | Epicardial cell subclusters C1 and C2 | Key transcription factor in epicardial and coronary vessel development (Wu et al 2013, Developmental Biology). |
| Fibronectin (FN1) | Epicardial cell subclusters C1-C5 | Extracellular matrix protein that is well known to promote cell adhesion and migration during cardiac development (Benesh et al 2013, Molecular Biology of the Cell). Found to be enriched in subepicardium of fetal chick hearts (Tidball et al 1992, Anatomy and Embryology) |
| Periostin (POSTN) | Epicardial cell subclusters C2-C5 | Extracellular matrix protein that supports cell migration and adhesion (Snider et al 2009, Circulation Research) |
| Agrin (AGRN) | Epicardial cell subcluster C2, C3, and C5 | Glycoprotein implicated in inducing cardiomyocyte proliferation and dedifferentiation (Bassat et al 2017, Nature) |
| **Myocardial Lineage** | | |
| Myoglobin (MB) | Cardiomyocyte subcluster C2 | Carries and stores oxygen within muscle tissues including cardiac (Garry et al 2003, Trends in Cardiovascular Medicine) |
| Paired Like Homeodomain 2 (PITX2) | Cardiomyocyte subcluster C1 | Transcription factor involved in directing asymmetric morphogenesis during cardiogenesis (Franco et al 2003, Trends in Cardiovascular Medicine) |
| Cardiac troponin C (TNNC1) | Cardiomyocyte subcluster C1-C3 | Cardiac type and slow skeletal troponin subunit |

**Supplemental Table 3: Additional gene markers for thymosin beta-4 enriched cell subclusters (Section 4)**

| **Gene name (Symbol)** | **Cell type(s)** | **Details and references** |
| --- | --- | --- |
| myosin heavy chain 15  (MYH15) | TMSB4X High Cells Subcluster 1 | Ortholog of MYH6 and a well-known ventricular cardiomyocyte marker. |
| Iroquois Homeobox 6 (IRX6) | TMSB4X High Cells Subcluster 1 | Transcription factor that was found to be localized to ventricular endocardium in fetal mouse hearts (Mummenhoff et al 2001, Mechanisms of Development) |
| Actin alpha cardiac muscle 1 (ACTC1) | TMSB4X High Cells Subcluster 1 | Cardiomyocyte specific actin |

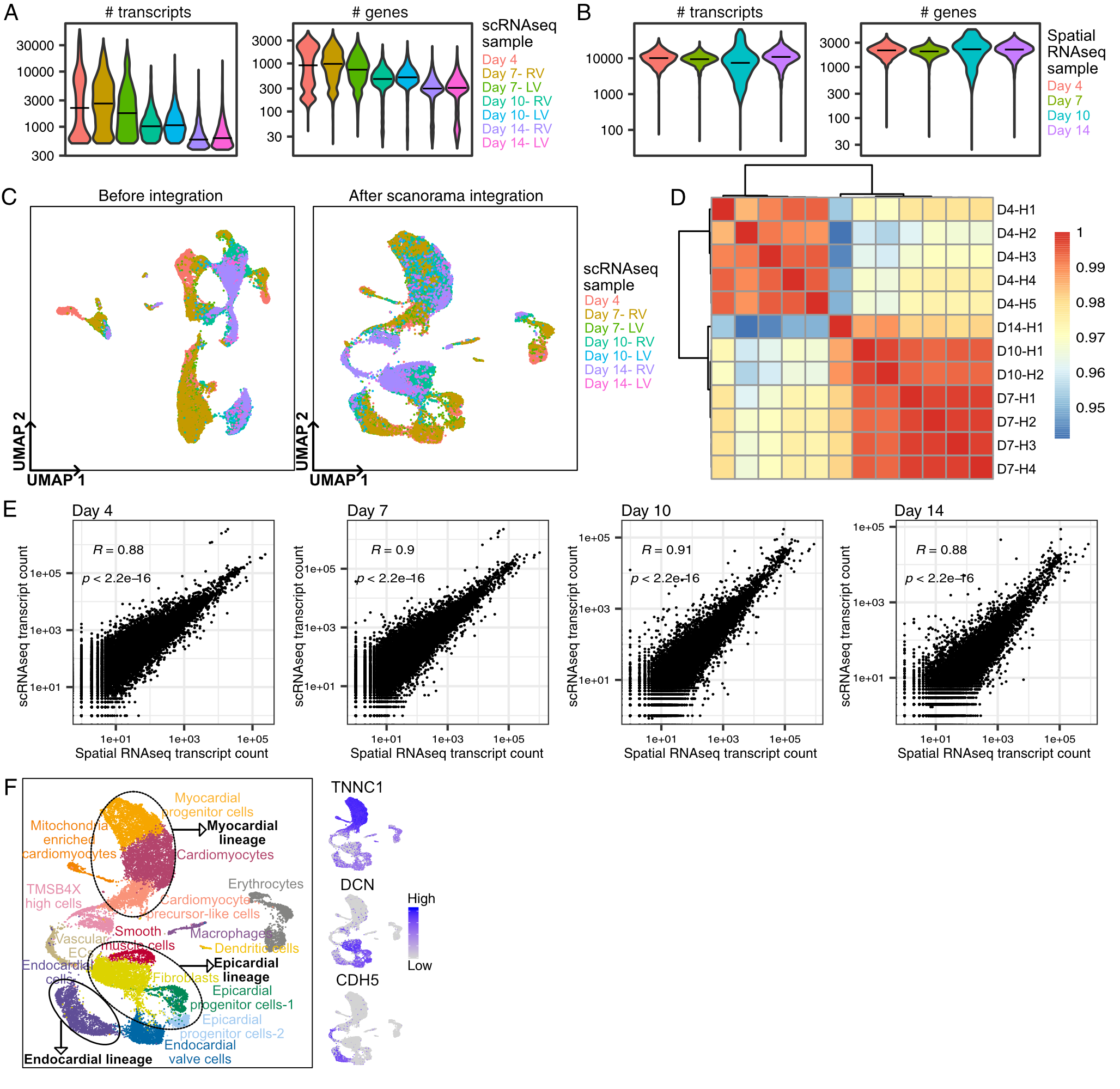

**Supplementary Figure 1: Time matched scRNA-seq and spatial RNA-seq atlas of developing fetal chicken hearts. A)** Number of unique transcripts per cell (left) and number of unique genes detected per cell (right) in seven scRNA-seq samples across four developmental stages. **B)** Number of unique transcripts per spot (left) and number of unique genes detected per spot (right) in four spatial RNA-seq samples across four developmental stages. **C)** UMAP projection of 22,315 scRNA-seq cells color labeled by scRNA-seq samples before (left) and after (right) scanorama based integration and batch effect removal. **D)** Pearson correlation between total spot gene expression in individual hearts (12 hearts) across 4 developmental stages in spatial RNA-seq data. Gene expression in hearts within the same developmental stage correlated well with each other (Pearson; R > 0.97). **E)** Pearson correlation between total gene expression in scRNA-seq samples and spatial RNA-seq samples from the same development stage (left to right: day 4 to day 14 stages). **F)** UMAP projection of 22,315 scRNA-seq cells clustered by gene expression and color labelled by cell types with black circles highlighting the endocardial, epicardial, myocardial lineages (left). Feature plots showing expression of lineage specific markers: TNNC1 for myocardial, DCN for epicardial, and CDH5 for endocardial (right).

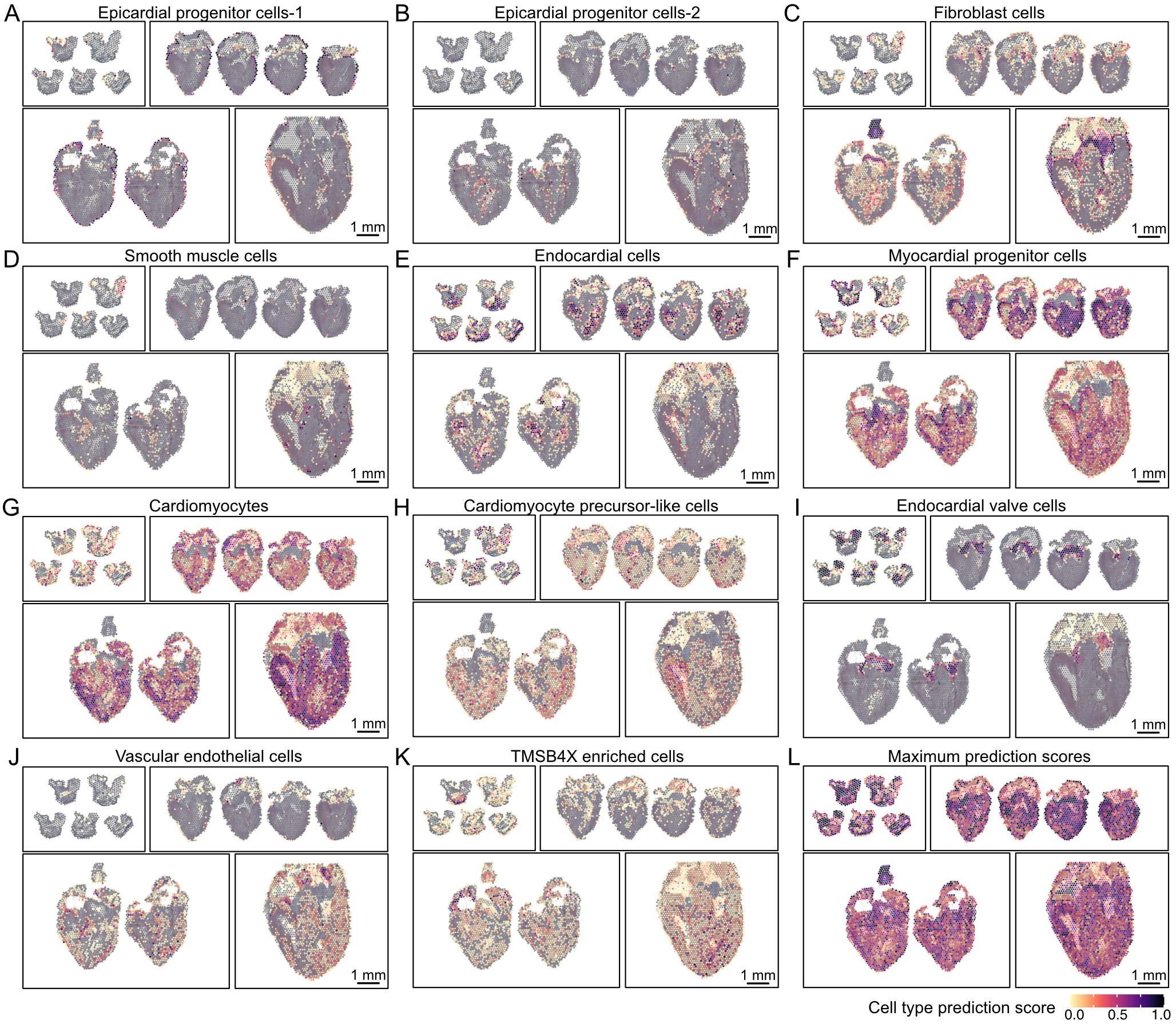

**Supplementary Figure 2: Anchor based integration of scRNA-seq and spatial RNA-seq and label transfer A-K)** Spatial cell type prediction scores derived by transferring scRNA-seq cell types on spatial RNA-seq data using an anchor based integration and batch correction method (Seurat-v3). Only Visium spots located under tissue are shown. **A)** Epicardial progenitor cells- 1. **B)** Epicardial progenitor cells- 2. **C)** Fibroblast cells. **D)** Vascular smooth muscle cells. **E)** Endocardial cells. **F)** Myocardial progenitor cells. **G)** Cardiomyocytes. **H)** Cardiomyocyte precursor cells. **I)** Endocardial valve cells. **J)** Vascular endothelial cells. **K)** TMSB4X high cells. **L)** Maximum cell-type prediction score for spatial RNA-seq spots.

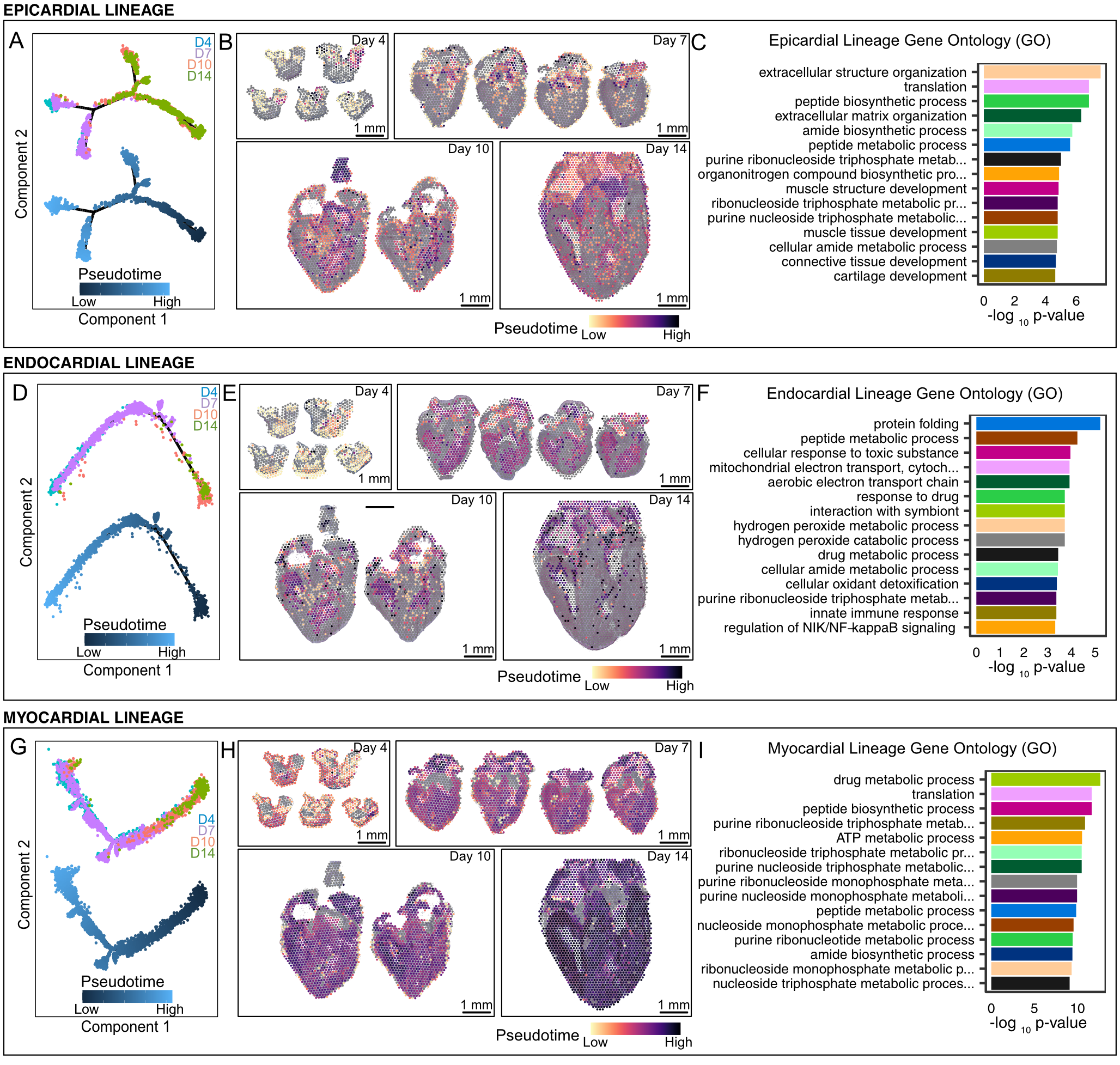

**Supplementary Figure 3: Monocle based lineage analysis and gene ontology for cardiac lineages. A, D, G)** Monocle v2 DDR-Tree projection of scRNA-seq cells from individual cardiac lineages labelled by developmental stage (top) and monocle pseudotime (bottom). **B, E, H)** Spatially resolved spatial RNA-seq spot pseudotime (monocle- based) for cardiac lineage across developmental stages. Spot pseudotime was estimated using a similarity map between scRNA-seq cells and spatial RNA-seq spots. **C, F, I)** Top 15 gene ontology (GO) terms for genes significantly correlated with pseudotime. Significant genes for gene ontology (GO) analysis were selected using differential marker analysis between sub-clusters within lineages with p-value threshold < 10^-15^. **A, B, C)** Epicardial lineage **D, E, F)** Endocardial lineage **G, H, I)** Myocardial lineage.

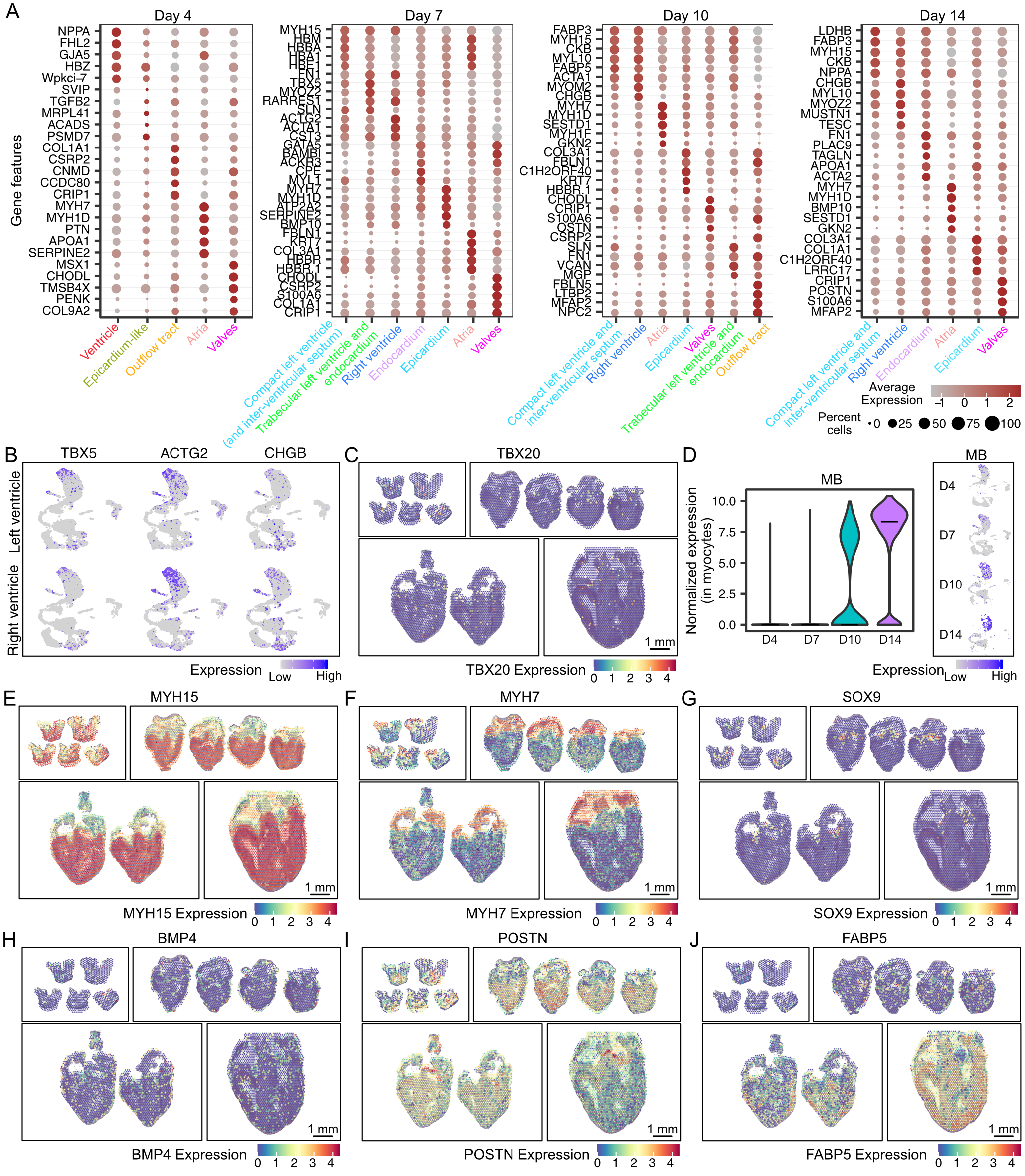
**Supplementary Figure 4: Spatial RNA-seq reveals spatially restricted genes in cardiac tissue during development. A)** Gene expression of region specific differentially expressed genes in spatial RNA-seq data. Regions were labelled by clustering of gene expression for barcoded spots in spatial RNA-seq data. Size of the dot represents the percent of spots in the spot clusters expressing the gene marker and the color intensity represents the average expression of the gene markers in that spot cluster. **B)** Feature plots showing the normalized scRNA-seq expression of TBX5, TBX20, and ACTG2 genes in cells from left and right cardiac ventricles. **C)** Spatially resolved normalized gene expression for TBX20 across all four developmental stages. **D)** Violin plot showing the distribution of normalized scRNA-seq expression of Myoglobin (MB) in myocardial lineage across developmental stages. Insets show normalized MB expression across stages in the entire scRNA-seq dataset. **E-G)** Spatially resolved normalized gene expression for spatially restricted genes across all four developmental stages. **E)** MYH15 significantly upregulated in ventricular tissue. **F)** MYH7 expression mostly restricted to atrial tissue. **G)** SOX9 expression restricted to valve tissue. **H-J)** Spatially resolved normalized gene expression for spatially restricted genes in specific developmental stages. **H)** BMP4 overexpressed in the outer lining of epicardium on day 7 and day 10. **I)** POSTN expressed in outer epicardium lining and myocardium on day 7 and day 10. **J)** FABP5 expressed in vascular bundles present in compact myocardium on day 10 and day 14.

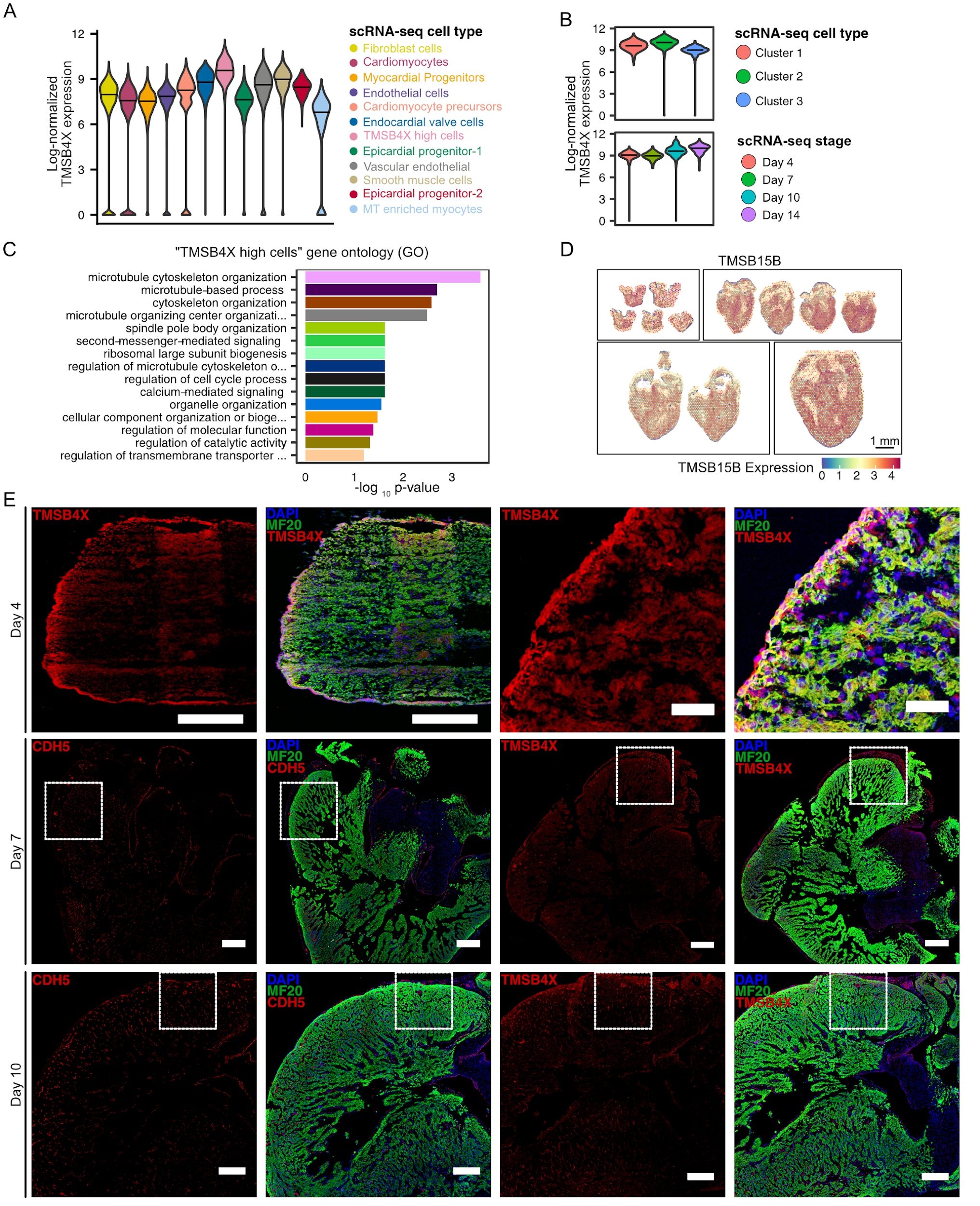

**Supplementary Figure 5: Characterization and validation of cell types in TMSB4X high cluster. A)** Log-normalized gene expression of Thymosin beta-4 (TMSB4X) in scRNA-seq cell type clusters across four developmental stages. **B)** Log-normalized gene expression of Thymosin beta-4 (TMSB4X) in subclusters within the “TMSB4X high cells” scRNA-seq cell cluster. TMSB4X expression is labeled by subcluster (top) and developmental stage (bottom). TMSB4X expression increases with developmental stage with TMSB4X high cells cluster. **C)** Top 15 gene ontology (GO) terms for genes significantly enriched in TMSB4X high cells scRNA-seq cluster. Significant genes for gene ontology (GO) analysis were selected with p-value threshold < 10^-15^. **D)** Spatially resolved thymosin beta 15B (TMSB15B) normalized gene expression across stages in chicken heart spatial RNA-seq data. **E)** Immunostaining images of chicken day 4 whole ventricles for cardiomyocyte cell marker MF20 (green) and thymosin beta-4 TMSB4X (red) at low magnification (Scale bar = 200µm) (top row, left) and high magnification (Scale bar = 50µm) (top row, right). Also high magnification tilescan immunostaining images of chicken whole heart sister sections of day 7 stage (middle row) and day 10 stage (bottom row) labeled for cardiomyocyte cell marker MF20 (green), and either endothelial cell marker CDH5 (red), or thymosin beta-4 TMSB4X (red) (Scale bar = 200µm). White boxes indicate regions of the ventricle that were magnified in the main Figure 2.
